# Molecular interactions of Chd8 in mouse brain highlights a role in chromatin-associated RNA processing

**DOI:** 10.64898/2025.12.11.693549

**Authors:** Emily Smith, Cesar P. Canales, Mathew W. Kenaston, Nicolas Seban, Ayanna A. Wade, Ethan Fenton, Cory Ardekani, Stephanie Lozano, Nickolas C. Chu, Daniela Perla, Ksenia Medvedeva, Kaitlyn McCafferty, Yingjing Guo, Aspen Kopley-Smith, Abigail Ignasiak, Dylann Cullinane, Karol Cichewicz, Gabriela Grigorean, Priya S. Shah, Alex S. Nord

## Abstract

The chromatin remodeler *CHD8* is a model risk gene for neurodevelopmental disorders (NDDs). While CHD8 has nucleosome remodeling capacity, evidence suggests it participates in processes beyond chromatin regulation, raising questions about function in the brain and role in NDDs. We defined CHD8 interactions using a comprehensive multimodal omics approach. Immunoprecipitation followed by mass spectrometry (IP-MS) identified a complex interaction network enriched for chromatin remodeling, RNA processing, and cytoskeletal proteins in neonatal mouse forebrain. We implemented CHD8-TurboID in HEK293T cells, validating IP-MS signatures and further identifying a role in mitosis. In addition to DNA, Chd8 complexed with RNA in mouse forebrain, with affinity for genes associated with RNA splicing and nervous system development and overlap between Chd8-bound mRNA and promoters. Finally, Chd8 interaction affinity with RNA splicing factors was reduced in *Chd8* haploinsufficient mice. These findings expand understanding of CHD8 function and identify a dosage-sensitive NDD-relevant role in chromatin-associated RNA processing.

## INTRODUCTION

Autism spectrum disorder (ASD) and neurodevelopmental disorder (NDD) risk genes are strongly enriched for chromatin remodeling factors (CRFs)^1,23^, a surprising link considering the broad expression and essential nature of many of these genes^4–6^. CRFs are ATP-dependent proteins that control chromatin dynamics via regulating nucleosome spacing and cooperating with transcription factors and histone modifiers to control transcription^7^. ASD/NDD-associated CRFs are expressed across many cell types and have pleiotropic functions and essential roles in development and cancer^8^. Heterozygous loss-of-function and missense mutations for CRFs cause ASD/NDDs, implicating haploinsufficiency as the relevant disease state. As such, complete loss-of-function studies may identify essential CRF functions, but fail to have ASD/NDD relevance. A major challenge in understanding CRF-associated molecular mechanisms driving ASD/NDDs is determining the spectrum of functional interactions of these pleiotropic proteins in the brain and which of these functions are disrupted by haploinsufficiency.

The CRF *CHD8* (Chromatin Helicase DNA-binding protein 8) is among the most significant ASD/NDD risk genes from rare variant studes^9–11^. *CHD8* mutation carriers display core ASD symptoms as well as macrocephaly, intellectual disability, sleep disorders, and gastrointestinal issues^10,12^. *CHD8* mutations have also been reported in schizophrenia and obsessive compulsive disorder^9,10^. Functional studies have further nominated CHD8 as a master regulator of neurodevelopmental ASD-associated pathways^9^ via contribution to expression control of other ASD risk genes^13^. *CHD8* haploinsufficiency impacts processes such as neuron differentiation, synapse development, axon guidance, and cell adhesion^14^.

*Chd8*^+/-^ mice exhibit macrocephaly, perturbations to neuronal development and signaling, and behavioral and cognitive phenotypes^15–17^. While a number of studies have shown reduced CHD8 expression leads to transcriptional phenotypes, functional studies in the brain have primarily focused on the phenotypic outcomes of haploinsufficiency rather than underlying molecular mechanisms. Thus, the direct mechanistic link between *CHD8* haploinsufficiency and ASD/NDDs remains poorly understood.

*CHD8* belongs to the CHD gene family^18–20^, which is comprised of nine proteins and divided into three subfamilies based on structural homology and function^21,22^. Along with *CHD8*, *CHD1*, *CHD2*, *CHD3*, *CHD4*, *CHD5* and *CHD7* are high-confidence NDD genes^1,23–31^. All CHD proteins are characterized by N-terminal tandem chromodomains and a central SNF2-like ATPase domain^32^. The chromodomain is involved in remodeling chromatin structure^22,32–34^. Functional analyses have demonstrated that CHD chromodomains mediate chromatin interactions via directly binding to DNA, RNA and methylated histone H3^32,35,36^. CHD8 is a member of the third subfamily of CHD proteins that includes CHD5-9 and are the least studied members of the CHD family^21,22,24,32^ characterized by additional domains in the C- terminal region, including tandem Brahma and Kismet (BRK) domains^22,32^. In the nucleus, CHD8 localizes to promoters and enhancers^37,38^ and in vitro assays show CHD8 has the capacity to shift nucleosomes^39^. Thus, a majority of studies have pursued the hypothesis that the primary ASD/NDD- relevant activity of CHD8 is as a chromatin remodeler^40–43^. Comparisons of chromatin accessibility have revealed large scale changes in homozygous knockout or knockdown *CHD8* models^44–46^. However, several in vitro studies showing pronounced chromatin changes were conducted in homozygous *CHD8* knockout models, whereas heterozygous models generally exhibited more modest or non-significant effects^44,45,47^. Other functions of CHD8 may be sensitive to single-copy loss-of-function mutations linked to ASD and related NDDs.

Despite significant attention on *CHD8* as a model ASD/NDD-linked CRF, the specific functions of CHD8 in the brain remain known, representing a major barrier toward understanding the role of CHD8 in the brain and CHD8-associated and CRF-associated mechanisms underlying NDDs. Unbiased Interaction studies can give a comprehensive picture of protein function and recent studies have mapped protein- protein interactions (PPIs) of high-risk ASD genes^48,49^, but none focus specifically on the CHD8 interaction network or were performed using brain tissues. Beyond the value of such studies for understanding CHD8, there is hope that single gene studies in neural systems will identify convergent molecular mechanisms across ASD/NDD associated genes, for example shared interaction networks or molecular activities. Towards addressing this, we performed Chd8 affinity purification mass spectrometry^50^ in wild-type and mutant *Chd8^5bpdel/+^*neonatal mouse brain and orthogonal in vitro proximity labelling^51,52^ in HEK293T cells and analyzed Chd8 interactions with mRNA and DNA in mouse brain. Our results capture a complex molecular interactome of Chd8 and highlight functions beyond chromatin remodeling in the brain, in particular involvement in chromatin-associated co-transcriptional RNA processing, that may be susceptible to *Chd8* haploinsufficiency.

## RESULTS

### Affinity mass spectrometry defines the Chd8 protein interactome in mouse forebrain

CHD8 is composed of three main protein domains, along with two disordered regions. These domains include an N-terminal tandem chromodomain, a SNF2-like ATPase helicase, and two C-terminal BRK domains^19,53^ (Figure 1A). Anti-CHD8 antibodies that target either the evolutionarily conserved C- or N- terminus are commercially available and have been used in previous studies of Chd8 expression and interactions. We compared the performance of four anti-Chd8 antibodies targeting N-terminus (human AA 325-350) or C-terminus (human AA 2252-2302) of CHD8 via western blot analysis of postnatal day 2 (PND2) forebrain extracts from wild-type and heterozygous mutant *Chd8* mice carrying a 5bp frameshift deletion in exon 5^15^ (Chd8*^5pdel/+^*) (Figure S1). Both the N-terminus and C-terminus antibodies identified a protein of about 280kDa, the size corresponding to the full-length of Chd8, with a significant decrease of expression for this band in *Chd8^5bpdel/+^*mouse brain lysates. Both antibodies also recognized other products that do not correspond to the size of any described isoform of Chd8 and that do not show abundance variability in *Chd8^5bpdel/+^* mice, indicating some non-specific binding. (Figure S1). C-terminal antibodies appeared to be more specific with only one additional band with a size of 30kDa, while N-terminal antibodies resulted in an additional 18 or 22 bands, with sizes ranging from 22kDa to 240kDa. These results show that both C- and N-terminus antibodies detect full-length Chd8 protein, though with antibody-specific off-target interactions.

**Figure 1.**
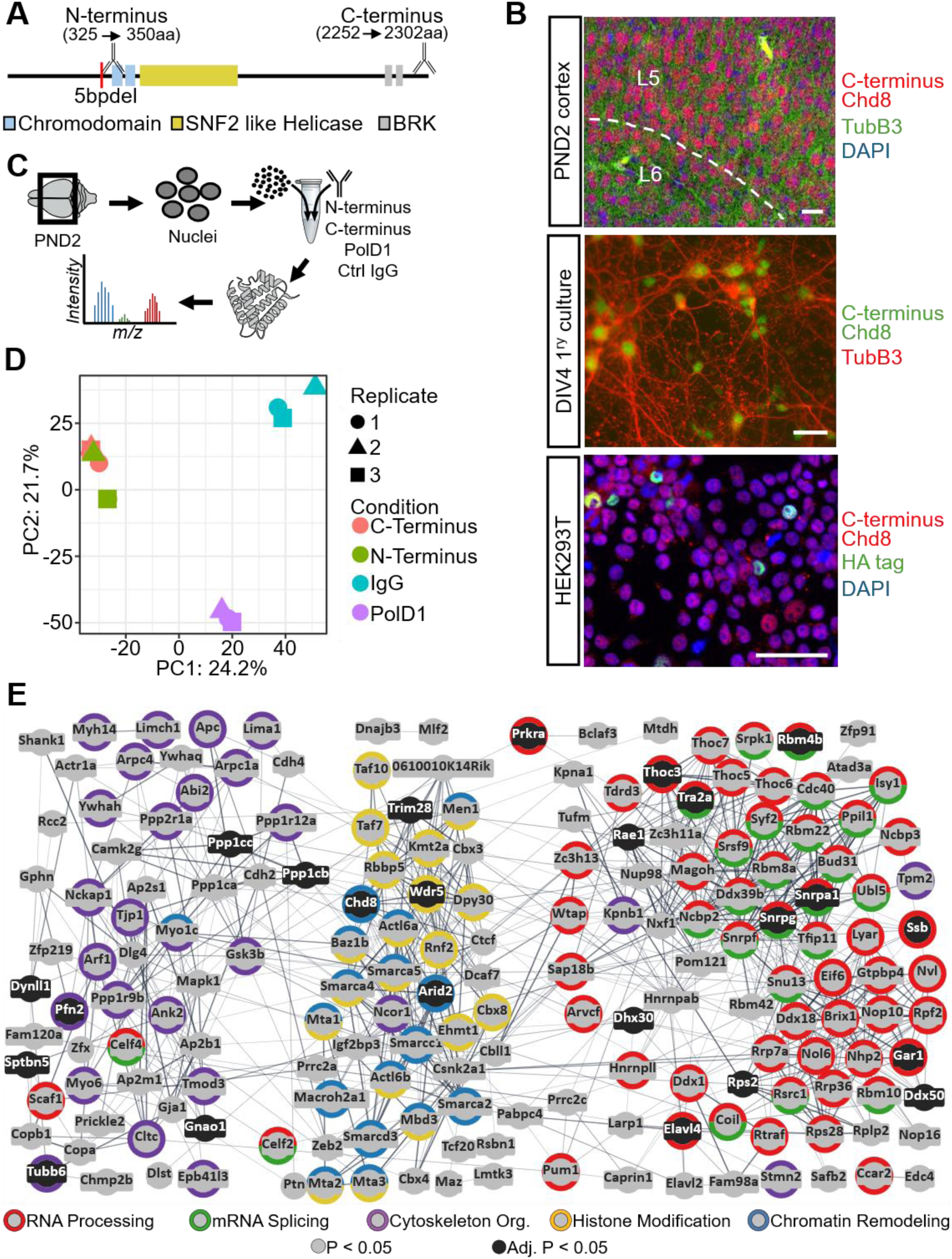
Chd8 interaction in PND2 mouse forebrain is enriched in chromatin remodeling and RNA processing proteins. **A)** Schematic of Chd8 protein domains with location of antibody epitopes indicated. The red line corresponds to the location of the 5bp deletion mutation to generate the *Chd8^5bpdel/+^* mouse line. **B)** Immunohistochemistry of Chd8 localization in PND2 mouse cortex, DIV14 primary neuron culture and HEK293T cells. Scale bar = 50µm. **C)** Schematic of endogenous immunoprecipitation approach. **D)** PCA plot showing peptide spectra variance of PC1 and PC2 in the four IP conditions, dimension reduction was performed on all proteins detected after IP. **E)** STRING Chd8 PPI network of all proteins considered significant for both Chd8 antibodies by SAINTexpress and DEP. Color of the outside ring indicates the annotated GO term. Inner circle color indicates the level of significance calculated via DEP. Network was made using default settings for a full STRING network indicating both functional and physical protein associations.

Chd8 has predominantly nuclear localization, though a recent study reports extra-nuclear CHD8^54,55^. We assessed the subcellular localization of Chd8 using immunocytochemistry (ICC) and immunohistochemistry (IHC) using the more specific C-terminus Chd8 antibody. We validated nuclear localization across multiple systems: the mouse cortical plate at PND2, primary mixed neuronal cultures in vitro, and immortalized HEK293T cells transfected with a CHD8-HA overexpression construct as an alternative to anti-Chd8 antibody methods. In all systems, Chd8 signal primarily colocalized with the nuclear marker DAPI, confirming its nuclear enrichment (Figure 1B). Some perinuclear localization was observed in the cultured cells including the primary mixed neuronal cultures and the CHD8-HA expressing HEK293T cells, with the latter potentially due to ectopic expression. (Figure S1). Cell fractionation was also performed on PND2 wildtype mouse forebrain. Cytoplasmic, nuclear, and whole cell fractions were collected and analyzed by western blot, using both the N-terminal and C-terminal Chd8 antibodies. While some Chd8 protein was detected in the cytoplasmic cell fraction, the vast majority of Chd8 was in the nuclear fraction, confirming strong bias for nuclear enrichment in the brain (Figure S1). In summary, our findings reinforce that Chd8 is primarily nuclear localized in the neonatal mouse brain overall and in neurons specifically, with cytoplasmic or perinuclear localization likely reflecting context-dependent activity or in vitro artifacts.

To characterize the protein interaction landscape of Chd8, we performed endogenous immunoprecipitation (IP) using nuclear-enriched protein lysates derived from PND2 mouse forebrain followed by liquid chromatography-tandem mass spectrometry (LC-MS/MS) (Figure 1C). Considering issues with anti-Chd8 antibody specificity, we used both C- and N-terminus antibodies and an intersection strategy to enrich for true interactions. IP was carried out on native lysates using anti-Chd8 antibodies, with anti-Pold1 antibody, targeting a nuclear DNA polymerase protein associated with DNA repair^56^ as a nuclear comparison, and IgG as a negative control. While Chd8 and Pold1 are likely to partially overlap in interactions, we include Pold1 to test for specificity of Chd8 interactions. Three biological replicates were performed for each condition. Peptides were identified and mapped to proteins via Spectronaut (see Methods). Western blotting of the IP before LC-MS/MS was performed to validate Chd8 enrichment (Figure S2). A short C-terminus Chd8 isoform has been described in previous publications^19,57^, however, neither Western blot nor IP-MS indicated presence of such a protein (Figures S1 and S2). Principal component analysis of LC/MS-MS spectral data after IP of all bait proteins showed separation between Pold1 and IgG, while the N-terminal and C-terminal Chd8 antibodies clustered together (Figure 1D). Together, these results demonstrate robust and reproducible recovery of Chd8-specific interaction partners, validating our IP-MS approach for defining the Chd8 interactome.

To identify protein-protein interactions, we compared spectral values from each antibody to IgG control using two complementary statistical methods: Significance Analysis of INTeractome Express (SAINTexpress)^58,59^ and Differential Enrichment analysis of Proteomics data (DEP)^60,61^ (Figure S2). Chd8 itself was significantly enriched by both methods and both antibodies (Figure S2). SAINTexpress identified 571 candidate interactors for the N-terminal Chd8 antibody and 508 for the C-terminal antibody. Intersecting these sets yielded 358 shared proteins considered putative Chd8 interactors. DEP analysis P < 0.05 identified 559 and 434 interactors for the N-terminal and C-terminal antibody, respectively, with 352 shared between them. At a more stringent cutoff of DEP adjusted P < 0.05, there were 48 candidate interactors for the N-terminal Chd8 antibody and 55 for the C-terminal antibody, with 25 shared hits. Intersecting proteins identified by both antibodies via SAINTexpress and DEP analysis P < 0.05 produced a high confidence network of 222 protein set representing robust and reproducible Chd8-associated interactions while retaining potential weaker partners (Table S1). The 222 proteins of the Chd8 interaction network are enriched for ASD genes^62^, including 41 genes on the Simons Foundation Autism Research Initiative (SFARI) gene list, with 17 categorized as high-confidence ASD genes (permutation test, P=0.048, Figure S3). Next, we applied the same workflow to identify proteins interacting with Pold1 (Figure S3, Table S1). Compared to the Pold1 interaction set, 56/222 at P-value < 0.05 and 11/24 at Adjusted P-value < 0.05 were specific to the Chd8 IP-MS networks, indicating a largely distinct Chd8-specific network. Consistent with this, the overlap between interactions identified between Chd8 C- and N-terminus antibodies was significantly higher than between either Chd8 and Pold1 antibodies for all comparisons (Figure S3). Collectively, these results reveal a reproducible and specific Chd8 IP-MS interactome that is strongly enriched for ASD risk genes, underscoring the biological relevance of Chd8-associated protein complexes in neurodevelopment.

We used STRING to visualize the Chd8 protein interaction network and identify enriched gene ontology (GO) terms using the set of 222 Chd8 interactions (Figure 1E, Table S2). As expected for a chromatin remodeler, proteins involved in histone modification and chromatin organization were prominently enriched in the Chd8 interaction network. Notably, we also observed enrichment of proteins associated with RNA processing, mRNA splicing, and cytoskeleton organization. While the two Chd8 antibodies strongly overlapped in functional enrichment of the identified interactions, the C-terminus antibody showed stronger affinity for RNA processing associated proteins, while the N-terminus had higher affinity for chromatin associated proteins (Figure S3). With the exception of the cytoskeleton category, major GO terms from the full 222 set of Chd8 interactions were also enriched in the 56 Chd8-exclusive interactors not identified in Pold1 IP-MS (Figure S4, Table S2). In contrast, functional enrichment of Pold1-exclusive interactions captured different terms, with strongest enrichment for Cytoskeletal organization and the Nuclear pore complex, with support in the literature for these Pold1 interactions^63^ (Figure S4, Table S2). Of the 24 Chd8 interactions that passed DEP Adjusted P-value < 0.05 for both Chd8 antibodies, twelve were annotated as having RNA binding function, three with Chromatin remodeling, and five were associated with Cytoskeleton. Two recent publications that performed IP of CHD8 in iPSCs also found strong enrichment of proteins associated with mRNA processing^46,49^. At the individual protein level, one protein was shared between all three CHD8 studies, and 10 proteins were shared between our mouse brain interaction network and the larger iPSC interaction network^49^. These overlaps were greater than expected by chance (permutation test, P=0.0003 and P=0.0009, respectively, Figure S4). Overall, our IP-MS results capture a complex network of Chd8 protein interactions in mouse brain and identify functions of Chd8 in the nucleus beyond chromatin remodeling.

### Chd8 interacts with chromatin remodeling, RNA processing, and cytoskeletal complexes

To explore the finer organization within the full network, we applied Markov Cluster (MCL) analysis. MCL is an unsupervised, cluster algorithm that utilizes the concept of random walk to calculate the probability of staying in one cluster or going outside to a less related node when “walking” along a weighted edge network. This identifies clusters of functionally related proteins inside the larger network of all Chd8 protein interactors. We expanded this analysis to the 926 proteins that were identified using either Chd8 antibody via either SAINTexpress or DEP to capture other constituents of protein complexes of interest associated with the 222 prioritized interaction partners (Figure 2A). MCL clustering using a granularity parameter of six produced clusters that generally belonged to the same protein complex or were known interactors of the complex, with cluster sizes ranging between 37 and 2 proteins (Figure S5). The spliceosome and proteins associated with the spliceosome was the largest identified cluster with 37 total proteins including eleven proteins from the 222 prioritized set. Another example of interaction with RNA processing machinery was a cluster of 6 proteins from the exon junction complex. The second largest cluster included proteins associated with the ribosome. Multiple chromatin remodeling protein complexes were identified, including the NuRD, MLL, SWI/SNF complexes. All three of these complexes included five proteins from the restricted set. For cytoskeleton organization, the two largest clusters with the most proteins from the restricted set include the AP2 adaptor complex and the Arp2/3 complex. These results indicate that Chd8 interacts with multiple distinct complexes, linking chromatin remodeling, RNA processing, and cytoskeletal regulation.

**Figure 2.**
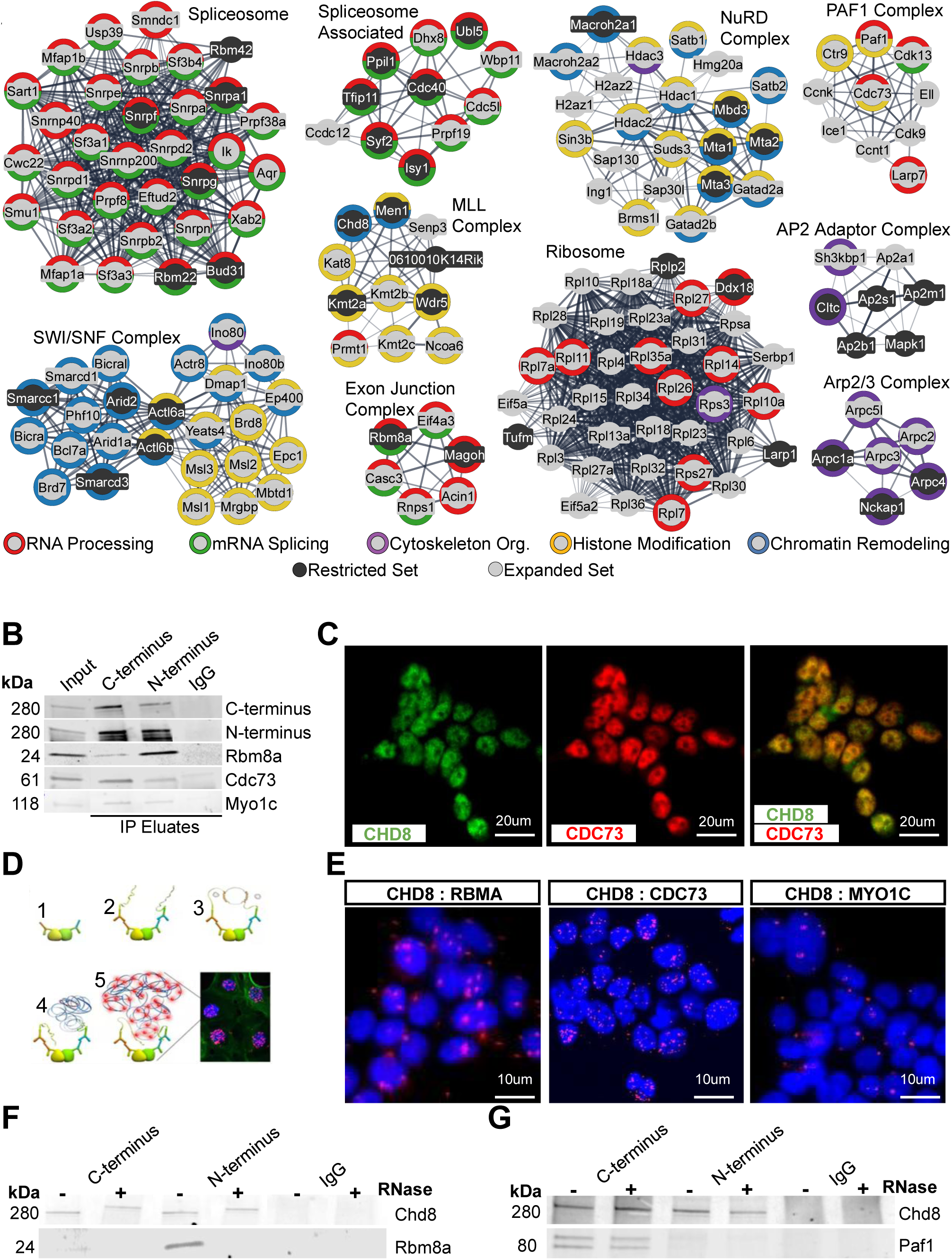
Chd8 interacts with complexes associated with mRNA splicing, chromatin remodeling, and cytoskeleton organization. **A)** MCL clustering of STRING Chd8 protein-protein network with all proteins identified by either Chd8 antibody via either SAINTexpress or DEP at a P-value < 0.05 significance cutoff. MCL clustering was performed with the granularity parameter set to 6. Color of the outside ring indicates the annotated GO term. Inner circle color indicates if the protein was included in the more stringent 222 Chd8 interacting protein set. Network was made using default settings for a physical STRING network indicating the proteins are part of a physical complex. **B)** Western blotting of Co-IP validation of Chd8 protein interactions. Co-IP was performed using either the C-terminus or N- terminus Chd8 antibody, indicated by the top labels. The antibody used for western blotting is indicated to the right and the molecular weight (kDa) of the band to the left. **C)** Colocalization validation of protein interactions. **D)** Schematic of Proximity Ligation Assay (PLA) approach. PLA visualizes protein interactions via fluorescent signal amplification when targets are in close molecular proximity. **E)** PLA validation of Chd8 protein interactions. Nuclei were stained blue with Hoechst, with red dots indicating locations where the two target proteins are interacting. **F-G)** Western blotting of Chd8 Co-IP after RNase treatment. +/- indicates the presence or absence of RNase, respectively. The top labels indicate the antibody used for Co-IP. **F)** Co-IP of Rbm8a. **G)** Co-IP of Paf1.

We validated select candidate interactors from the Chd8 protein network using multiple methods. Candidate proteins for confirmation were selected based on antibody compatibility and relevance. First, we performed co-immunoprecipitation (Co-IP) (Figure 2B). Nuclear lysates from PND2 mouse forebrain were subjected to IP using either the N-terminus or C-terminus Chd8 antibody, with IgG serving as an isotype control. Among the interactors, Rbm8a, a component of the exon junction complex involved in mRNA splicing that remains with mature transcripts during cytoplasm export^64–66^, was robustly detected in both Co-IP conditions, with stronger enrichment observed using the N-terminus antibody. We validated interactions with the PAF1 complex (PAF1C), a key regulator of RNA Polymerase II elongation and chromatin structure^67^. While Cdc73 was the only PAF1C member identified as significant in both Chd8 antibody pulldowns relative to IgG, all core components of the complex, including Paf1 and Leo1, were recovered in at least one IP. Western blotting confirmed successful pulldown of Cdc73 with both antibodies, with higher efficiency observed for the C-terminus antibody. Finally, we validated interaction with Myo1c, a member of the unconventional myosin protein family known to function as an actin-based molecular motor^68^ . Both the N-terminus and C-terminus Chd8 antibodies pulled down Myo1c with comparable efficiency. Although primarily cytosolic, one isoform of Myo1c localizes to the nucleus and interacts with RNA Polymerase I and II during transcription initiation^69^. This interaction is also consistent with a study reporting CHD8 association with microtubules and cytoskeletal elements^54^. Other interactions validated via Co-IP included Paf1, Leo1, and Ddx39b, with all targets successfully recovered by both Chd8 antibodies (Figure S6).

We next validated interactions for these same candidates using orthogonal immunocytochemistry (ICC) and proximity ligation assays (PLA) in human HEK293T cells (Figure 2C-E, Figure S6). As an example, we confirmed nuclear co-localization of CHD8 and the PAF1C (Figure 2C), with localization of CDC73 and LEO1 as a positive control and localization of CHD8 and MAP2, a cytoplasmic protein involved in microtubule stabilization, as a negative control (Figure S6). All candidates tested with PLA showed expected interactions. Moderate PLA signal in the nucleus was observed for the CHD8–RBM8A and CHD8–CDC73 pairs (Figure 2D-E). For both co-localization and PLA assays, a CDC73-LEO1 pair used as a positive control for proteins interacting in the nucleus and a CHD8-MAP2 pair was used as a negative control for proteins localized to different subcellular compartments (Figure S6). Together, the Co-IP and PLA experiments validate specific Chd8 physical interactions in the brain with components of RNA processing, transcriptional control, and cytoskeletal machinery, supporting a multifunctional role for Chd8 in the nucleus. Lastly, considering the extensive interactions between Chd8 and RNA-associated proteins, we tested if candidate interactions were dependent on RNA by treating PND2 mouse brain lysate with RNase before IP. Rbm8a was successfully co-immunoprecipitated under untreated conditions with both antibodies but was undetectable after RNase treatment (Figure 2F). In contrast, Paf1 remained detectable after RNase treatment (Figure 2G). These results suggest that some Chd8 interactions are dependent on RNA, further highlighting a role for Chd8 linking promoter-associated chromatin and transcriptional processes with co-transcriptional RNA processing.

### Development of CHD8-TurboID and deployment in human HEK293T cells identifies similar proteins as endogenous IP-MS in mouse brain

Affinity-based IP-MS can be impacted by technical considerations associated with IP pulldown and antibody performance. Considering limitations of anti-Chd8 antibodies and broad interaction networks from IP-MS, we sought to identify Chd8 interactions using an orthogonal approach. For this, we implemented TurboID, a biotinylation-based proximity labeling method and applied this to map CHD8 proximity-based interactions in human HEK293T cells. We generated a fusion protein (CHD8-Turbo) by attaching an HA-tagged TurboID to the N-terminus of CHD8 under the control of a CMV promoter (Figure 3A). This construct was transiently expressed in HEK293T cells to assess subcellular localization and biotinylation activity (Figure 3B). As expected, CHD8-Turbo localized predominantly to the nucleus. Limited cytoplasmic and perinuclear localization was also observed, potentially due to overexpression or artifacts from ectopic expression. Biotinylation activity was detected only in cells transfected with CHD8-Turbo and only after treatment with exogenous biotin. Notably, biotinylation signals colocalized entirely with the TurboID fusion protein, confirming construct-specific activity.

**Figure 3.**
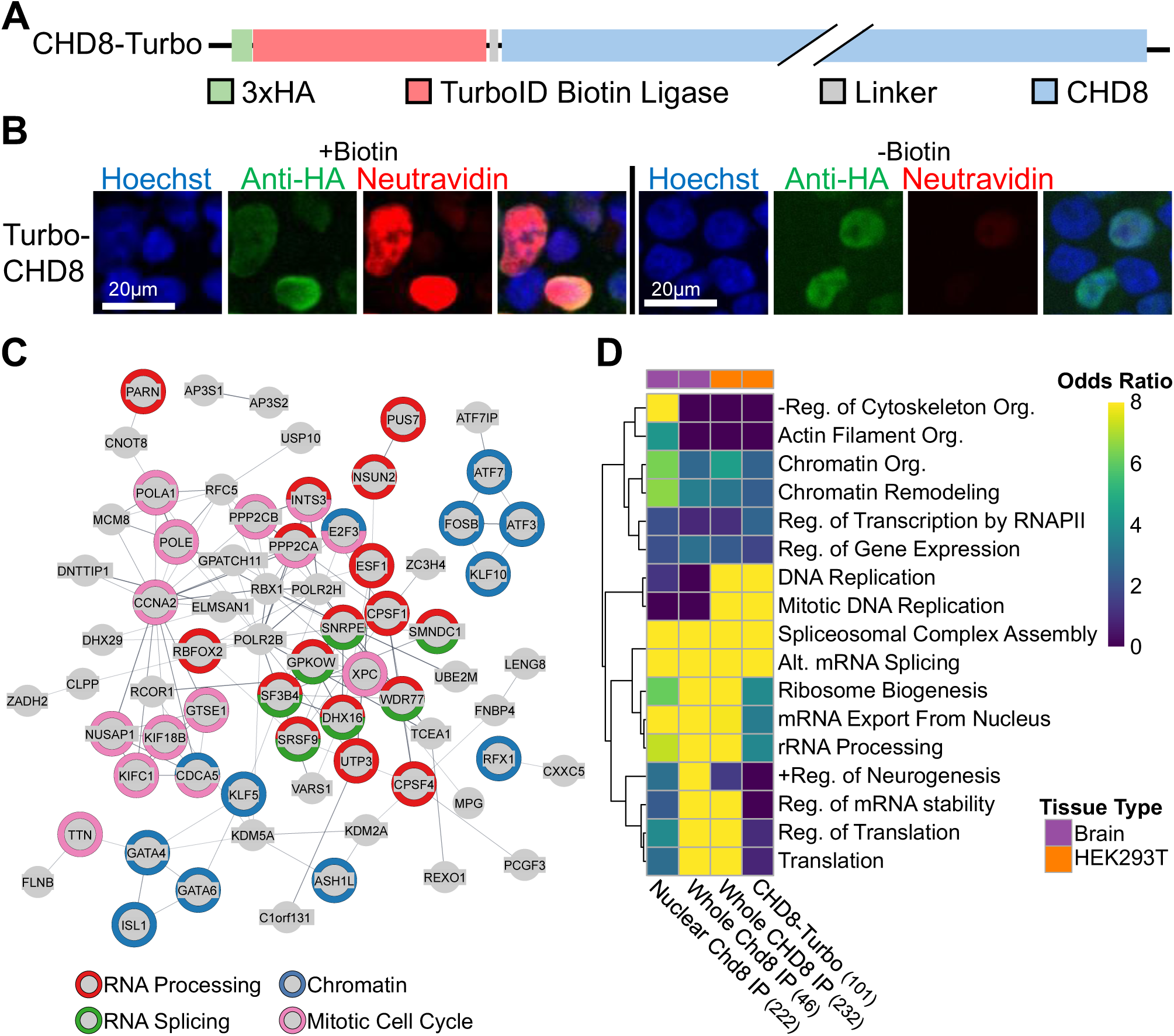
CHD8 targeted TurboID in HEK293T cells as a parallel approach to endogenous immunoprecipitation. **A.** Schematic of the protein construct used for TurboID. **B)** Epifluorescence images validation of localization and biotinylation activity of CHD8-Turbo in HEK293T cells. Protein localization was stained by anti-HA antibodies and biotinylation activity was stained by fluorescently conjugated Neutravidin. **C)** STRING PPI network of the CHD8-Turbo fusion protein in HEK293T cells. Color of the outside ring indicates the annotated GO term. Network was made using default settings for a full STRING network indicating both functional and physical protein associations. **D)** Heatmap of GO term enrichment across multiple Chd8 pulldown conditions. The number of proteins included in each condition is indicated in parentheses.

To identify CHD8-associated proteins in HEK293T cells, we performed TurboID-based proximity labeling followed by LC-MS/MS. CHD8-Turbo was transiently expressed in HEK293T cells, and twenty- four hours post transfection, cells were incubated with biotin-containing media (CHD8-Turbo +Biotin) or regular media (CHD8-Turbo -Biotin). Biotinylated proteins were affinity purified using NeutrAvidin- conjugated magnetic beads. To distinguish specific CHD8 interactions we applied the same strategy as for IP-MS comparing CHD8-Turbo +Biotin to CHD8-Turbo -Biotin. Using SAINTexpress, 997 proteins were identified as significant relative to CHD8-Turbo -Biotin. Using DEP, there were 1299 interactors at P-value < 0.05 and 148 at Adjusted P-value < 0.05 (Table S3). The large number of significant hits demonstrates sensitivity of the Chd8-Turbo system between biotin and no biotin conditions and indicates CHD8 comes into proximity with a large number of nuclear proteins. To prioritize strong interactions, significant proteins identified by SAINTexpress and DEP at Adjusted P-value < 0.05 were intersected, producing a final set of 101 proximity-based candidate interactions used for further analysis (Figure S7, Table S3). The HEK293T CHD8 TurboID interaction network was enriched in similar GO terms to those found in the IP-MS mouse forebrain Chd8 network, with RNA processing and RNA splicing as top hits and Chromatin associated proteins also strongly significant (Figure 3C, Table S2). Cytoskeleton Organization, which was enriched in the mouse brain dataset, was not enriched in the TurboID dataset. Conversely, the TurboID set was enriched in Mitotic Cell Cycle. At the individual protein level, the CHD8 interaction networks from mouse brain (222 proteins) and TurboID in HEK239T cells (101 proteins) shared only three proteins, which may be due to detection method or biological system. When expanded to proteins identified by either SAINTexpress or DEP at P-value < 0.05, the IP-MS and TurboID CHD8 interaction networks share 313 proteins (permutation test, P=0.0003, Figure S7). Thus, CHD8-TurboID identified overlapping biological processes but relatively distinct specific proteins interactions for CHD8 versus IP-MS.

We assessed concordance of results across endogenous immunoprecipitation from nuclear-enriched or whole cell lysates of PND2 mouse forebrain, endogenous immunoprecipitation from whole-cell lysate of HEK293T cells, and CHD8- targeted TurboID proximity labeling in HEK293T cells. For IP-MS datasets, we used the interactors identified by SAINTexpress and reached P-value < 0.05 with DEP for both C- and N-terminus antibodies. For the Chd8-TurboID, we used the set that were identified by SAINTexpress and reached Adjusted P-value < 0.05 with DEP. All conditions showed strong enrichment for multiple RNA processing/splicing-related GO terms as well as Chromatin-associated GO terms at somewhat lower levels (Table S4). GO terms related to Mitosis and DNA replication were prominently enriched in both IP-MS and TurboID HEK293T datasets but not in brain datasets, likely reflecting differences in proliferative activity between PND2 mouse brain and HEK293T cultures. The only condition showing enrichment for actin cytoskeleton organization and negative regulation of cytoskeleton organization was the nuclear-enriched Chd8 IP-MS from mouse brain. Differences in GO term enrichment between PND2 forebrain conditions, where chromatin-associated proteins were under- represented in the whole cell data, may be attributed to both cell fractionation and the mass spectrometry methods, as data-dependent acquisition (DDA), which captures most abundant peptides, was used for the whole cell lysates versus data-independent acquisition (DIA), which provides more comprehensive coverage, was used for the nuclear-enriched condition. Despite these differences, there was strong concordance across all CHD8 interaction proteomics experiments, supporting a consistent role for CHD8 in nuclear RNA processing alongside its well-established function in chromatin remodeling, as well as context-dependent interaction with proteins involved in cytoskeletal and mitotic processes.

### Chd8 preferentially interacts with mRNA of genes associated with RNA splicing and other general cellular processes and nervous system development

Given the enrichment of RNA-associated proteins in the Chd8 protein network and RNA-dependent results from Co-IP experiments, we hypothesized Chd8 may bind RNA or participate in ribonucleoprotein complexes, and we tested this via IP (Figure 4). We isolated RNA following IP of Chd8 from PND2 mouse forebrain tissue. IPs were performed using both N-terminus and C-terminus Chd8 antibodies, with Hnrnpa2b1, a known RNA-binding protein and putative Chd8 protein interactor, H3K4me3, a promoter-associated modified histone protein, and IgG as controls (Figure 4A). As expected, Hnrnpa2b1 IP yielded the highest RNA recovery, while IgG showed little to no RNA enrichment. Both Chd8 antibodies pulled down RNA, with the N-terminus antibody recovering RNA at levels comparable to Hnrnpa2b1, and the C-terminal antibody pulling down approximately half as much (Figure 4B). To verify nucleic acid composition, IP eluates were treated with DNase or RNase prior to quantification (Figure 4C). DNase treatment had minimal impact on nucleic acid levels in all conditions except H3K4me3, where a modest reduction suggested partial DNA association consistent with its promoter localization. In contrast, RNase treatment significantly reduced nucleic acid levels across all conditions except IgG. These results demonstrate that Chd8 directly or indirectly associates with RNA, supporting the hypothesis that Chd8 participates in ribonucleoprotein complexes and may influence co-transcriptional processes.

**Figure 4.**
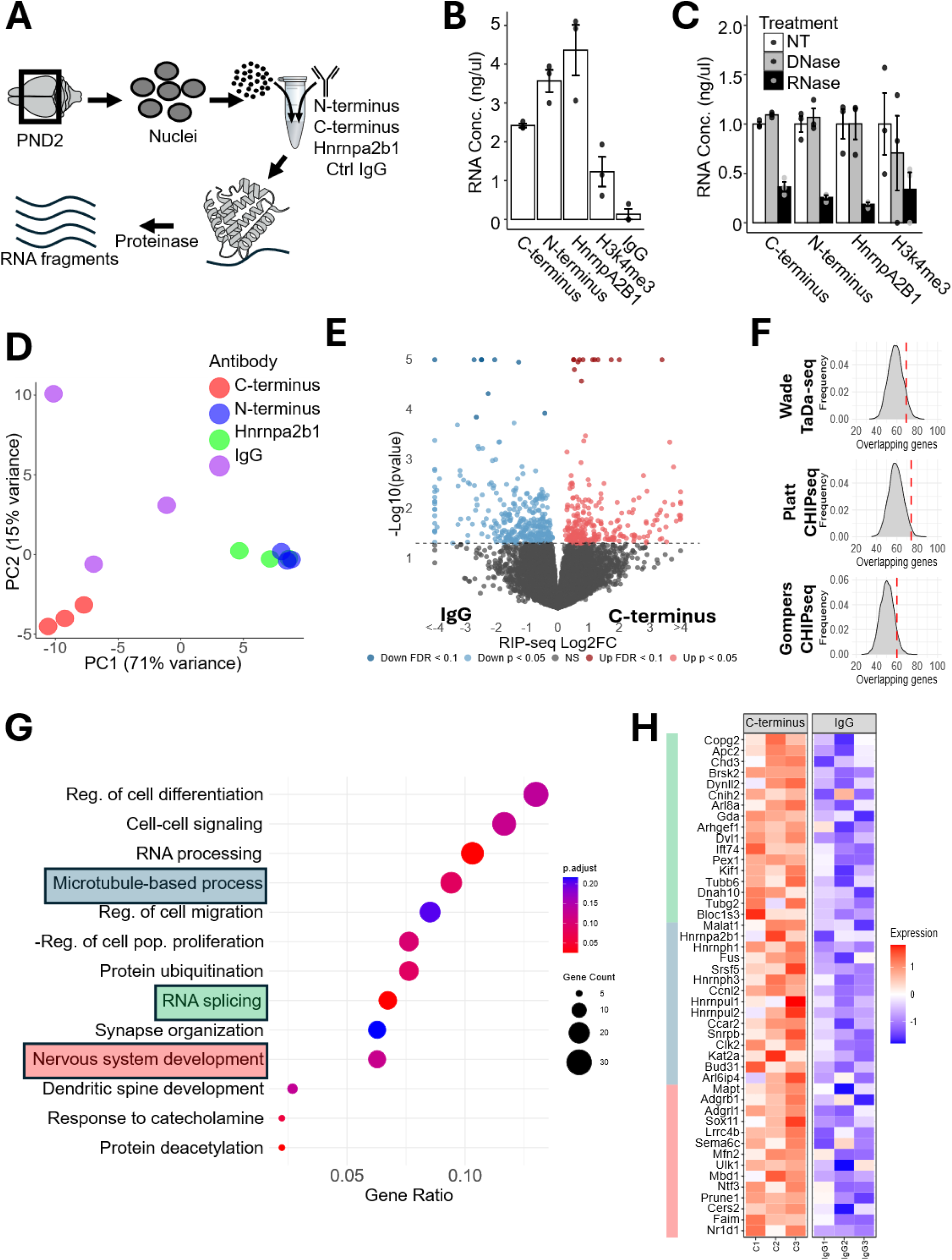
P2 Chd8 RIP-seq and RNA-seq identifies RNA binding partners of Chd8 and cellular deficits during forebrain development. **A)** Schematic of PND2 forebrain RIP-seq method. **B)** Quantification of RNA that was isolated from Chd8 Co-IP from PND2 mouse forebrain. **C)** Quantification of RNA that was isolated from Chd8 Co-IP after DNase and RNase treatment. **D)** PCA plot showing transcriptional variance of PC1 and PC2 in the four IP conditions used for RIP-seq, dimension reduction was performed on the top 500 variable genes. Antibodies used for IP conditions are indicated by color. **E)** Volcano plot showing log fold change and significance of mRNA enriched after IP via the C-terminus antibody vs IgG control. Significant enrichment in C-terminus vs IgG is shown in red (P value < 0.05) with highly significant enrichment (FDR < 0.1) shown in dark red. 263 genes showed significant enrichment in the C-terminus RIP-seq vs IgG. **F)** Density plots showing the frequency of gene overlaps from permutation tests comparing the top 1000 C-terminus RIP-seq enriched genes and Chd8-bound genes from, the top 1000 Chd8 TaDa-seq (Wade 2021) genes (P = 0.0874), the top 1000 Chd8 CHIP-seq (Platt 2017) genes (P = 0.0232), and the 856 Chd8 CHIP-seq (Gompers 2016) genes (P = 0.0784). Observed intersections in shown in red. **G)** Gene ontology enrichment of genes significantly (P < 0.05) enriched in C-terminus RIP-seq. Strength and significance of enrichment, as well as number of DEGs present in gene set is indicated. Gene sets with DEGs highlighted in panel H are colored. **H)** Heatmap showing the z-scale normalized counts of DEGs present in indicated gene sets across C-terminus and IgG IP samples. Color to the left indicates the enriched gene set.

Having shown that Chd8 Co-IP yields RNA, we adapted a protocol for RNA Immunoprecipitation followed by sequencing (RIP-seq) to identify transcripts that were enriched in Chd8 interactions (see Methods, Figure 4). Again, using mouse PND2 forebrain lysates, we performed IP using both the N- and C-terminus Chd8 antibodies as well as Hnrnpa2b1 and IgG on three independent biological replicates. Principle components analysis of RIP-seq count data showed separation between IgG and the C-terminus Chd8 antibody, while the N-terminus Chd8 antibody and Hnrnpa2b1 clustered together (Figure 4D). As the C-terminus antibody showed greater specificity for Chd8 in initial testing and was more distinct relative to Hnrnpa2b1, we focused differential comparison on the C-terminus and IgG datasets, identifying 220 RNA transcripts that were significantly enriched via DESeq2 (P < 0.05) following Chd8-C pulldown (Figure 4E, Table S5). These results indicate that Chd8 associates with a defined subset of RNA transcripts in the developing forebrain, consistent with selective and biologically meaningful RNA-binding activity.

To test if Chd8 RIP-seq RNA interactions are associated with Chd8 genomic targets, we compared RIP-seq enriched transcripts with three published datasets mapping Chd8 DNA binding, two using antibody-dependent Chromatin Immunoprecipitation followed by sequencing (ChIP-seq) using adult mouse cortex^14,70^, and the other based on antibody-independent Targeted DamID followed by Sequencing (TaDa-seq) using E17.5 mouse cortex^71^. We used the top 1000 hits from these genomic binding datasets for comparison. The intersection of genes with promoter interactions from ChIP-seq and TaDa-seq and genes with the 220 transcripts enriched in RIP-seq at P < 0.05 was not significantly greater versus random permutation. However, when extended to the top 1000 RIP-seq enriched transcripts, there was modest evidence for overlap for both ChIP-seq and Tada-seq identified Chd8 genomic targets (permutation test, ChIP-seq P=0.0232 and P=0.0784, and Tada-seq P=0.0874, Figure 4F). We next tested Chd8 RIP-seq hits for functional enrichment, yielding a number of significant GO terms including general cellular functions (e.g., cell differentiation and RNA processing), as well as brain-specific processes (e.g., nervous system development and synapse organization) (Figure 4G). Enrichment patterns for individual target mRNA were reproducible across replicates and specific to the C-terminus Chd8 antibody compared to Hnrnpa2b1, as shown by the sets of Chd8 RIP-seq enriched transcripts annotated to Microtubule-based process, RNA splicing, and Nervous system development (Figure 4H). Overall, the RIP-seq results indicate direct or indirect interactions between Chd8 and RNA in the nucleus and support link between CHD8 promoter-interactions and RNA transcription and processing.

### *Chd8* haploinsufficiency reduces affinity of RNA splicing protein interactions in mouse forebrain

Heterozygous mutation to *CHD8* is associated with NDD and ASD and increased brain size. Complete loss causes embryonic lethality in mice and conditional forebrain knockout leads to failure of the cortex to develop^72,73^. This differing dosage effect on the brain may be due to different impacts between complete loss of function and haploinsufficiency on Chd8 function. To test whether Chd8 haploinsufficiency affects affinity of protein interactions, we performed LC-MS/MS following IP with either the N-terminus or C-terminus Chd8 antibodies in the forebrain of heterozygous mutant *Chd8^5bpdel/+^* mice. Of the 926 proteins that were detected by either Chd8 antibody, 162 proteins passed a P-value < 0.05 significance cutoff, with 109 proteins increased in *Chd8^+/+^* mice and 53 increased in *Chd8^5bpdel/+^*(Figure 5A, Table S6). 22 of the proteins increased in *Chd8^+/+^*forebrain also passed an Adjusted P-value < 0.05 significance cutoff. No proteins increased in *Chd8^5bpdel/+^*passed the Adjusted P-value significance cutoff. Proteins that were significantly increased in *Chd8^+/+^* forebrain were enriched for mRNA processing, mRNA splicing, and spliceosomal assembly (Figure 5B, Table S6). Proteins that were significantly increased in *Chd8^5bpdel/+^*forebrain were enriched in proteins associated with actin filament organization, neuron projection development, and organelle organization. Compared to the background of all 926 proteins with evidence for Chd8 interaction, only RNA capping passed an FDR < 0.05 cutoff, for proteins significantly increased in *Chd8^+/+^*, with Actin filament organization in *Chd8^5bpdel/+^*.

**Figure 5.**
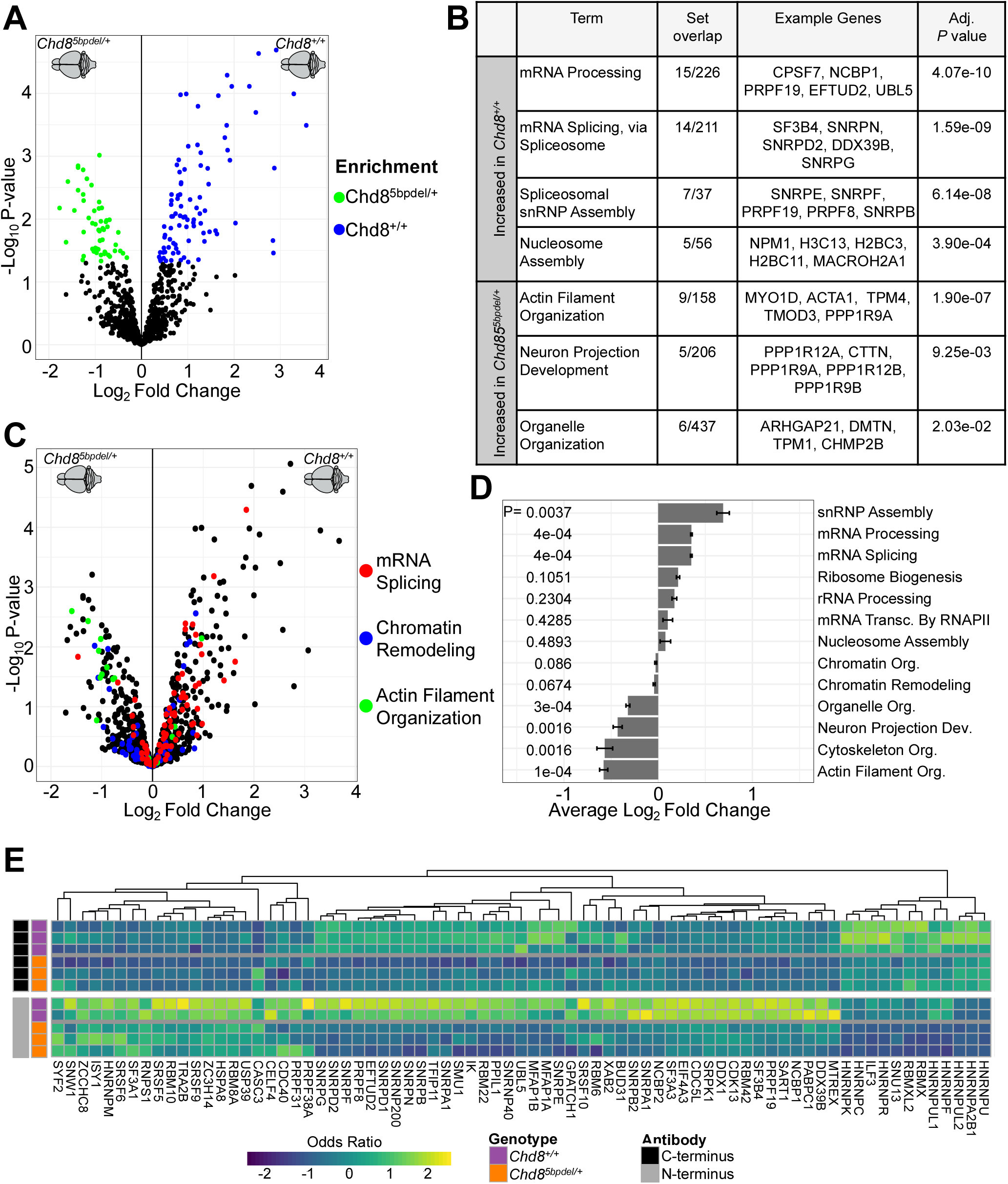
Interactions with mRNA splicing proteins are decreased in *Chd8^5bpdel/+^* mice compared to wildtype. **A)** Volcano plot of individual protein pulldown in *Chd8^5bpdel/+^*and wildtype mice including proteins identified by either Chd8 antibody via SAINTexpress or DEP at a P-value < 0.05 cutoff. Green and blue proteins are significantly differential for *Chd8^5bpdel/+^* or *Chd8^+/+^* forebrain at a significance of P-value < 0.05. **B)** GO terms enriched in proteins significantly increased in *Chd8^5bpdel/+^* or *Chd8^+/+^* forebrain. Set overlap indicates the number of significantly increased proteins and total detected indicates the total number from all proteins analyzed. **C)** Volcano plot of individual protein pulldown in *Chd8^5bpdel/+^*and wildtype mice. Color indicates the annotated GO term. **D)** The observed average fold change of multiple GO terms. SEM and p-values calculated by permutation test. Sample size for each permutation corresponds to the number of annotated proteins for each GO term. **E)** Heatmap comparing Odd Ratio values across Chd8 antibody and mouse genotype for all proteins associated with mRNA Splicing, via Spliceosome (GO:0000398).

Notably, 56 out of 79 proteins mapping to mRNA Splicing (GO:0000398) were increased in *Chd8^+/+^* forebrain (Figure 5C,E). To test for pathway-level impacts on interaction affinity, we calculated the average fold change of interactors mapping to each GO term and used a permutation framework to test if the GO term average logFC was significantly different from random sampled sets drawn from the full set of Chd8 interactors. Average logFC of mRNA splicing and mRNA processing interactors was significantly increased in *Chd8^+/+^*forebrain, while actin filament organization was significantly increased in *Chd8^5bpdel/+^* (Figure 5D). No pathway-level differences were observed for nucleosome assembly or chromatin-associated GO terms. These results suggest that Chd8 haploinsufficiency leads to selective reduction of Chd8 affinity for the set of proteins involved in RNA processing, while interactions related to chromatin remodeling remain unaffected and actin cytoskeleton interactions were slightly increased.

## DISCUSSION

The strong link between CRF haploinsufficiency and ASD/NDDs, with *CHD8* as a leading example, raises questions about the functions in the brain of these broadly expressed and frequently pleiotropic proteins, as well as how reduced expression impacts specific CRF activities. Much of the focus on Chd8 with respect to NDD-associated mechanisms has been on nucleosome remodeling and regulation of chromatin state. Our interaction findings using two orthogonal proteomics methods deployed on ex vivo neonatal mouse brain and in vitro human HEK293T cells capture broader functional roles, suggesting Chd8 not only regulates nucleosome positioning and chromatin state, but is also involved in RNA processing and in cytoskeletal interactions and mitosis. We further show that Chd8 directly or indirectly interacts with RNA and that Chd8-RNA interactions are enriched for specific biological processes and linked to Chd8 promoter binding. Finally, via proteomic comparison of *Chd8^5bpdel/+^* and wildtype littermates, we found suggestive evidence for decreased affinity between Chd8 and RNA processing proteins. Our findings expand understanding of the molecular interactome of Chd8 and identify potential NDD-relevant molecular functions, specifically highlighting chromatin-associated RNA processing.

The extensive and complex Chd8 protein interactome identified here is consistent with previous proteomics and functional studies and reported direct or indirect involvement of Chd8 in many biological processes. Recent studies have focused on its role in nucleosome remodeling, epigenetic state, and promoter-associated protein interactions^16,74–76^. Aligned with these studies, our results suggest that rather than a small number of strong interaction partners, Chd8 interacts with a broad variety of proteins and complexes. As expected, we observed strong enrichment for proteins and complexes associated with chromatin and epigenetic state (e.g., SWI/SNF, MLL, and NuRD) and with promoters and transcriptional control (e.g., RNA polymerase and PAF1 complexes). Many of the interactions identified here are related to processes and interactions reported for CHD8 that are involved in epigenetic and transcriptional regulation, including Wnt signaling^17,70^, cell cycle regulation^15,70,72^, cancer related processes and transcriptional regulation via interactions with β-catenin^18^, promoting transcription of E2F target genes^77^, recruiting histone H1 to negatively regulate p53^74^, and interacting with CHD7 to regulate RNA polymerase II^78^. Along with chromatin remodeling and epigenetic state, our results highlight Chd8 interactions with RNA and RNA processing as a central function of Chd8 in mouse brain and human cells. Previous studies have shown that CHD8 interacts directly with elongating RNA polymerase or interacts with other factors that promote RNA transcription^79,80^ and two recent in vitro IP-MS studies of CHD8 also found the strongest enrichment in interactions with RNA processing associated proteins^46,49^, evidence for the validity and relevance of these interactions. Extending beyond gene regulation, a recent study reported that CHD8 localizes to microtubules, and that its loss of function causes mitotic spindle defects, cell cycle stalling, DNA damage, and cell death^54^. Consistent with these findings, our interactome includes multiple cytoskeletal and mitotic regulators, including components of the actin-related Arp2/3 and AP2 adaptor complexes, that support association of CHD8 with the cytoskeleton and cell division machinery. Our results integrate previous candidate based and in vitro studies of Chd8 interactions, providing an unbiased and comprehensive perspective of the complex interactions and functions of this high-priority ASD/NDD-relevant protein. Overall, these findings reinforce a role of Chd8 in chromatin associated processes and highlight expanded roles in RNA processing and cytoskeletal interaction during neurodevelopment.

While any of the identified protein interactions and associated molecular functions may be impacted by *CHD8* haploinsufficiency, our results combined with recent studies highlight chromatin-associated co- transcriptional RNA processing as a candidate NDD-relevant and dosage-sensitive function. Genomic profiling studies have mapped widespread CHD8 binding across the genome, showing enrichment at promoter regions and strong association with genes involved in RNA processing, chromatin remodeling, and cell cycle regulation^77,81,82^. These findings, together with evidence for a role of CHD8 in nucleosome remodeling, have shaped a prevailing view that NDD pathology primarily arises from CHD8-dependent disruption of chromatin and epigenetic state. However, while homozygous Chd8 loss in mice or in in vitro knockout models causes marked perturbations in chromatin accesibility^83^, multiple studies have reported fairly minimal effects in heterozygous mutant mouse or knockdown in vitro models^15,70,84–86^ .

This suggests that in the context of haploinsufficiency, dysregulation of processes other than direct chromatin remodeling roles may contribute to pathophysiology. Supporting this, a recent study showed that while some human CHD8 variants linked to NDD cases impaired nucleosome remodeling in vitro, others that produced NDD-relevant phenotypes in mice did not impact nucleosome remodeling^39^, indicating that chromatin remodeling alone cannot explain CHD8-related pathology. Our results point to co-transcriptional RNA processing as an alternative NDD-associated mechanism impacted by Chd8 haploinsufficiency. We found that Chd8 directly interacts with RNA processing proteins and complexes with RNA, and that Chd8 affinity of RNA-associated protein interactions was reduced in *Chd8^5bpdel/+^* PND2 forebrain. Aligned with our results, another recent study identified altered histone modifications associated with RNA elongation in *CHD8* mutant cells and interaction between CHD8 and RNA Pol II complex and RNA associated proteins^46^. Specifically, these results align with our observation that CHD8 interacts directly with RNA-processing proteins and that CHD8-associated RNA interactions are reduced in mutant brain, together supporting a model in which CHD8 haploinsufficiency impacts neurodevelopment not only through enhancer and promoter regulation but also by altering co-transcriptional chromatin states required for proper RNA processing and splicing. These splicing defects previously described provide a relevant mechanistic backdrop for our findings, suggesting that disrupted chromatin-associated functions of CHD8 may impair recruitment or stabilization of RNA-processing complexes, thereby linking chromatin remodeling and RNA metabolism. Collectively, these studies expand the pathogenic paradigm for CHD8 beyond transcriptional regulation to include disrupted coupling between chromatin elongation dynamics and RNA-processing machinery as a contributing mechanism in NDD pathology. Moreover, several transcriptomic studies of *Chd8^+/-^* mice, as well as human iPSC and organoid models, have described altered RNA splicing, reinforcing the concept that perturbed RNA processing is a key molecular consequence of *CHD8* haploinsufficiency^15,86^. Furthermore, many annotated CRFs have been shown to interact with RNA, including the CHD family member CHD1, to be involved with co-transcriptional RNA processing^87,88^.

While Chd8 has no annotated RNA binding domain, it does contain a DEAH-box helicase domain^89–92^ which is involved in ATP-dependent RNA or DNA unwinding^93,94^. Chd8 could also participate in co-transcriptional protein-RNA complexes rather than functionally interacting with RNA itself. Future studies are needed to elucidate the role of CHD8 in the production of RNA or association with RNA-associated proteins during transcription, and to link any direct impacts on cellular RNA homeostasis with NDD-associated phenotypes.

Few studies have used both IP and proximity labelling approaches to identify PPIs, as we did here with endogenous anti-Chd8 antibodies and CHD8 TurboID, enabling the opportunity to compare results and providing orthogonal support to interactions. One study used both methods in combination to identify the protein interactome of a tumor-inducing effector in plants^95^, while two used them separately and compared identified proteins^51,96^. TurboID appears to be a more permissive approach, with a higher number of TurboID-identified proteins versus IP in these previous studies and our work in HEK293T cells. TurboID will biotinylate proteins in proximal environment of the bait, which may capture indirect and transient interactions that might not be pulled down by IP, and CHD8-Turbo -Biotin appears to serve as a cleaner control compared to IgG. Despite a larger number of identified interactions and similar GO terms among interactors in HEK293T cells and mouse brain, we found minimal overlap between individual proteins identified by Chd8 TurboID and IP. This mirrors findings for other chromatin associated proteins^96^ and there is evidence that TurboID and affinity-based IP have a preference for recovering distinct biological processes^51,96^. Nonetheless, the difference between IP-MS and TurboID raises some questions about how to prioritize and interpret results for individual interactions. Beyond differences between TurboID and IP, the CHD8-Turbo construct was also ectopically expressed at high levels. While further studies are needed to optimize Chd8 TurboID, development of this system overcomes antibody dependency and has the potential to address questions not tractable with Chd8 antibody IP, for example evaluating domain dependency of interactors or testing impact of patient CHD8 mutations on interaction affinity.

There are several limitations to this study. All commercially-available Chd8 antibodies had imperfect specificity (particularly Chd8 N-terminus antibodies). While use of knockout models can be used to identify off-target interactions, *Chd8* homozygous mutation is embryonic lethal in mice, so we were not able to use this strategy in this system. As an alternative, we used two different Chd8 antibodies and required significant detection in both. We further compared Chd8 and Pold1 interactions, with evidence for antibody specificity based on non-overlapping interactions. Finally, IP and MS methods influences interaction detection^97–99^, leading to possible biases from specific methods used here. With respect to CHD8 TurboID, we used full length CHD8 with TurboID fused to the N-terminus and ectopic expression levels that likely differ from endogenous CHD8 expression. In previous work, we showed fusion of a DNA Adenosine Methylation (DAM) domain to the Chd8 N-terminus did not impact CHD8 genomic interactions^100^, evidence that the CHD8-Turbo fusion should retain at least partial functionality, which is supported by general overlap between IP-MS and TurboID results. Finally, RIP-seq targeting Chd8 RNA interactions is a novel application and, as such, we lacked positive controls and note that the number of very strong Chd8-RNA interactions (FDR < 0.05) were few and there was overall modest enrichment over IgG, suggesting weak Chd8-RNA interaction preference and an extensive RNA interaction set. However, these patterns are relatively consistent with other RIP-seq studies on chromatin associated proteins^101,102^, and we show that Chd8 RIP-seq interactions differ from Hnrnpa2b1 and are linked to Chd8 genomic interactions. While these limitations should be considered, our results lay the foundation of future work characterizing the functional relevance of Chd8 interactions.

In summary, our study expands understanding of Chd8 molecular interactions in the mammalian brain and we developed a novel proximity labelling system that offers new avenues to characterize Chd8 protein function. Our results fill in gaps in understanding that are relevant to understanding CRF activity in the brain and the link between CRFs and ASD/NDDs, showing Chd8 has pleiotropic roles in neonatal mouse forebrain. Our findings highlight a role for Chd8 in co-transcriptional RNA processing based on the interactions with RNA-associated proteins, Chd8 IP yielding RNA, and links between Chd8- associated transcripts, promoters, and gene expression. Many NDD-associated genes have primary annotated functions related to RNA processing and recent studies of chromatin factors highlight interaction with RNA for many of these factors^44,72,103^, raising the possibility of convergence in ASD/NDD molecular pathology at the level of RNA processing. Overall, our results capture the complex nuclear interactome of Chd8 in vivo in the brain and highlight new directions of research on the role of Chd8 and other NDD-linked genes, including CRFs, in functions that impact the processing, transport, and stability of mRNA during brain development and in neuron function.

## ACKNOWLEDGMENTS

This work was supported by the National Institutes of Health National Institute of Mental Health (NIH NIMH) R01 MH120513 to A.S.N. and National Institutes of Health National Institute of Allergy and Infectious Disease (NIH NIAID) R01 AI170857 to P.S.S.. C.P.C. was supported by the NIH NIMH Autism Research Training Program fellowship (T32 MH073124-16). S.L. was supported by T32 GM 007377 and F31 HD113328-01A1. A.A.W. was supported by NIH NIMH F31 MH119789 and T32 GM007377.

## AUTHOR CONTRIBUTIONS

A.S.N. conceived the project with substantial contributions from E.S. and C.P.C.. E.S led the project execution, designed experimental work and oversaw most aspects of data generation and analysis. C.P.C. fully executed tissue collection and led the IHC, colocalization, and PLA experiments with contributions from D.P., K. Medvedeva, K. McCafferty, A.K-S., and A.I. E.S. fully executed all IP and pulldown experiments with contributions from A.A.W and Y.G. G.G. carried out LC-MS/MS and mass spectrometry data analysis. E.S., M.W.K., P.S.S., A.S.N. carried out proteomics data analysis. N.S. led most of the RNA-seq data analysis with contributions from K.C., E.F., C.A., S.A.L., N.C., and A.S.N. E.S., C.P.C, and A.S.N. wrote the manuscript.

## MATERIALS AND METHODS

### Whole cell protein extraction

Brains were extracted from 3 postnatal day 2 (PND2) mice (Table S7). The forebrain was collected then snap-frozen on dry ice for storage at -80℃. The forebrain tissue was suspended in 600μl lysis buffer (50mM Tris-HCl, 140mM NaCl, 10% Glycerol, 0.5% IGEPAL, 0.25% Triton X-100, protease inhibitor cocktail (Roche, 4693124001)) and manually homogenized. After a 30min incubation on ice, samples were sonicated with a probe sonicator (Qsonica CL-18, 20% amplitude, 10 cycles of 5sec on/off intervals) and placed back on ice to incubate for another 30min. Cell debris was collected by centrifugation (14000 g, 4°C, 10min) and the supernatant was moved to a new tube. Protein concentration was quantified using a BCA protein assay kit (Pierce, 23225) and samples were stored at -80℃ until use.

### Cell fractionation

Cell fractionation was performed using the NE-PER nuclear and cytoplasmic extraction kit according to manufacturer instructions (Thermo Scientific, 78835). Briefly, forebrain tissue was washed with PBS and manually homogenized in 600μl CERI supplemented with Halt protease and phosphatase inhibitors (Thermo Scientific, 78440). After a 10min incubation on ice, 60μl CERII was added and samples were incubated on ice for another one minute. Samples were centrifuged (16000g, 4℃, 5min) and the supernatant, containing cytoplasmic proteins, was collected. The nuclear pellet was resuspended in 300μl NER supplemented with Halt protease and phosphatase inhibitor. Nuclear samples were incubated on ice for 40min, vortexing every 10min, and centrifuged again (16000g, 4°C, 10min). The supernatant, containing all nuclear proteins, was collected. Both cytoplasmic and nuclear fractions were stored at -80℃ until use in western blot analysis or immunoprecipitation.

### Immunoprecipitation

Immunoprecipitation assays were performed using either HEK293T cells or postnatal day 2 (PND2) mouse forebrain tissue, in duplicate and triplicate, respectively (Table S7). Whole cell protein extracts from HEK293T cells and forebrain tissue were obtained following whole cell protein extraction. Nuclear protein extracts were obtained following cell fractionation, with each biological replicate consisting of pooled forebrain tissue from one male and one female PND2 mouse. For each immunoprecipitation reaction, 500μg of whole cell or 200μg of nuclear protein extract was diluted 1μg/1μl. For RNAse - treated samples, 3μg of RNase was added and samples were incubated for 30min at 37°C. All lysates were incubated with 20μl of a mixture of washed Protein A and Protein G Dynabeads (Invitrogen, 10001D and 10003D) on a nutator for one hour at 4°C. 8μg of primary antibody specific to the target protein was coupled to 40μl Protein a and Protein G Dynabeads mixture by incubation for 30min at 4°C on a nutator. CHD8 immunoprecipitation was performed with antibodies targeted to either the C- terminal (Bethyl, A301-225A) or N-terminal (Bethyl, A301-224A) end of CHD8. Other protein targets include MAP2 (Invitrogen, 13-1500), PAF1 (Bethyl, A300-172A), and POLD1 (Bethyl, A301-007A). For a general background control, 12μg of a rabbit IgG isotype control (02-6102, Invitrogen) was used. The precleared lysate was incubated with the antibody-coupled beads on a nutator overnight at 4°C. The antibody-bead complex was washed three times by gentle pipetting and nutation in ice-cold lysis buffer (50mM Tris-HCl, 140mM NaCl, 10% Glycerol, 0.5% IGEPAL, 0.25% Triton X-100, Roche protease inhibitor cocktail). The antibody-bead complex was either prepared for mass spectrometry or had protein complexes eluted for western blot analysis. For mass spectrometry, beads were washed three times in ice-cold 100mM TEAB (Thermo Scientific 90114) by gentle pipetting and nutation then resuspended in 100mM TEAB for submission to the UC Davis Proteomics Core. For elution, beads were resuspended in 60μl 6X Laemmli SDS buffer (375mm Tris-HCl, 9% SDS, 50% glycerol, 0.03% Bromophenol blue) and 5% β-mercaptoethanol and boiled at 70℃ for 10min. The supernatant was removed and used for western blot analysis.

### Generation of TurboID Constructs

The NLS-Turbo construct was provided by Alice Ting (Addgene plasmid #107171, http://n2t.net/addgene:107171, RRID:Addgene_107171). It has been previously characterized^52^ and no modifications to the plasmid were made.

The CHD8-Turbo construct was generated by modifying a CHD8 Human Tagged ORF Clone (OriGene, RG230753). The HA-tagged TurboID sequence from the NLS-Turbo construct was cloned by PCR amplification using CloneAmp HiFi polymerase (Takara, 639298). A linker sequence was added onto the 3’ end of the PCR fragment during amplification. The resulting PCR product was then gel purified (Thermo Scientific, K0692). The CHD8 ORF was digested using a standard restriction digest with enzyme AsiSI (NEB, R0630S). The gel purified TurboID PCR fragment and linearized CHD8 ORF clone were assembled by In-Fusion cloning (Takara, 638947), with the PCR fragment being inserted onto the 5’ end of the CHD8 sequence. Assembled plasmids were introduced by heat shock transformation into chemically competent *E. coli* cells (NEB, C3040H) and screened for correct insertion by whole plasmid sequencing verification (Genewiz, Plasmid-EZ). Once insertion of the HA-tagged TurboID sequence was verified, the GFP sequence was removed using a double restriction digest with enzymes NotI and PmeI (NEB, R0189S and R0560S). The plasmid was reassembled using In-Fusion, introduced into bacteria, and sequence verified as before.

### Characterization of TurboID Constructs

Poly-L-lysine (PLL) coated glass cover slips (Neuvitro, GG1815PLL) were placed into 12-well plates and quickly washed with sterile water. HEK293T cells were plated onto the coverslips with complete DMEM (Corning, 10013CV) supplemented with 10% fetal bovine serum (Gibco, 16140071). Cells were plated at 150,000 cells/well for transfection the next day. CHD8-Turbo and NLS-Turbo constructs were introduced by transfection using Lipofectamine 3000 (Invitrogen, L3000008) according to the manufacturer’s instructions in DMEM with 50% Opti-MEM (Gibco, 31985070). 24 hours after transfection, the media was replaced with warm complete DMEM containing biotin (Invitrogen, B1595) at a final concentration of 50μM. Cells were placed back at 37°C to incubate. After 15 minutes, cells were placed onto ice and washed 5 times with 1ml of ice-cold PBS (Gibco, 10010023). To fix, cells were treated with 4% paraformaldehyde at room temperature for 15 minutes. After treatment, cells were washed 3 times with PBS and stored at 4°C until use.

Transfected and non-transfected cells were permeabilized with 0.5% Triton-X100 (Sigma-Aldrich, X100) in PBS for 10 minutes at room temperature and blocked with 4% BSA (Thermo Scientific, J64655.22) and 1% normal goat serum (Cell Signaling, 5425S) in PBS with 0.1% Triton-X100 (PBST). Cells were labelled with anti-HA tag primary antibodies (1:500 dilution, Invitrogen, 26183) at 4°C overnight. The next day, cells were washed 3 times with PBST for 5 minutes at room temperature. An appropriate secondary antibody (AlexaFluor488, 1:1000 dilution, Jackson Laboratory, 715-545-151) was applied along with NeutrAvidin conjugated with Rhodamine Red-X (1:1000 dilution, Invitrogen, A6378) and Hoechst (1:10000 dilution, Invitrogen, H3570) for 1 hour at room temperature. After secondary staining, cells were washed 3 times with PBST for 5 minutes at room temperature and mounted on glass slides using Prolong Gold (Invitrogen, P36930). Images were recorded by epifluorescence using a Keyence BZ-X810 All-in-One fluorescence microscope.

### TurboID Protein Enrichment for Proteomics

HEK293T cells were plated and transfected as described above for characterization of constructs. Cells were transfected with either CHD8-Turbo or NLS-Turbo constructs (see Table S7 for replicate information). 24 hours after transfection, the media was replaced with warm complete DMEM containing biotin (Invitrogen, B1595) at a final concentration of 50μM, with half of the CHD8-Turbo transfected cells treated with DMEM without biotin. Cells were placed back at 37°C to incubate. After 30 minutes, cells were placed onto ice and washed 5 times with 1ml of ice-cold PBS. Cells were detached and collected via pipetting with 1ml of ice-cold PBS. Cells were centrifuged at 300g at 4°C for 3 minutes and supernatant was removed. RIPA lysis buffer (50 mM Tris, 150 mM NaCl, 0.1% SDS, 0.5% sodium deoxycholate, 1% Triton X-100) supplemented with 1x protease inhibitor cocktail (Roche, 4693124001) and 1mM PMSF (Thermo Scientific, 36978) was used to lyse cells. To ensure complete lysis, Cells were left on ice for at least 10 minutes on ice. Centrifugation at 13,000g at 4°C for 10 minutes to pellet cell debris and the supernatant was collected for further analysis. Protein concentration of the lysate was quantified using a BCA protein assay kit (Pierce, 23225). After quantification, lysates were stored at -80°C until further use.

Lysates from each condition, CHD8-Turbo +Biotin, CHD8-Turbo -Biotin, NLS-Turbo +Biotin, was prepared by diluting 300μg protein in 500μg RIPA buffer. For each sample, 25μl of streptavidin magnetic beads (Pierce, 88816) were washed twice with 1ml of RIPA lysis buffer. After washing, beads were incubated with the diluted protein lysates on a nutator overnight at 4°C. The next day, beads were collected with a magnetic rack and the supernatant was removed. Beads were washed sequentially as follows: twice with 1ml of RIPA buffer for 2 minutes, once with 1ml of 1M KCl for 2 minutes, once with 1ml of 0.1M Na2CO3 for 10 seconds, once with 1ml of 2M urea in 10mM Tris-HCl for 10 seconds, and twice with 1ml of RIPA buffer for 2 minutes. All washes were performed at room temperature. After washing, beads were prepared for LC-MS/MS by washing three times in ice-cold 100mM TEAB (Thermo Scientific 90114) by gentle pipetting and nutation, then resuspend in 100mM TEAB. Samples were submitted on-bead for LC-MS/MS analysis by the UC Davis Proteomics Core.

### Tryptic Digest and Liquid Chromatography

Protein samples on magnetic beads were washed four times with 200ul of 50mM Triethyl ammonium bicarbonate (TEAB) with a twenty minute shake time at 4C in between each wash. Roughly 2.5 ug of trypsin was added to the bead and TEAB mixture and the samples were digested over night at 800 rpm shake speed. After overnight digestion the supernatant was removed and the beads were washed once with enough 50mM ammonium bicarbonate to cover. After 20 minutes at a gentle shake the wash is removed and combined with the initial supernatant. The peptide extracts are reduced in volume by vacuum centrifugation and a small portion of the extract is used for fluorometric peptide quantification (Thermo scientific Pierce). One microgram of sample based on the fluorometric peptide assay was loaded for each LC-MS analysis.

Peptides were resolved on a Thermo Scientific Dionex UltiMate 3000 RSLC system using a PepSep analytical column (PepSep, Denmark): 150um x 8cm C18 column with 1.5 μm particle size (100 Å pores), preceded by a PepSep C18 guard column, and heated to 40 °C. Each injection is 0.6 μg of total peptide. Separation was performed in a total run time of 60 min with a flow rate of 500 μL/min, with mobile phases A: water/0.1% formic acid and B: 80%ACN/0.1% formic acid. Gradient elution was performed from 4% to 10% B over 4min, from 10% to 46% B over 44min, 46% to 99% B in 1.5min, down to 4% B in 0.5 min followed by equilibration for 10min.

### Mass Spectrometry and Data Analysis

#### Data Dependent Analysis

Data dependent analysis was used for the following immunoprecipitation samples: whole cell lysate from PND2 *Chd8^+/+^* mouse forebrain, whole cell lysate from HEK293T cells, and whole cell lysate from PND2 *Chd8^+/+^* mouse forebrain with RNase treatment. Peptides were analyzed on an Orbitrap Exploris 480 instrument (Thermo Fisher Scientific, Bremen, Germany), data acquired in data-dependent analysis mode. Spray voltage were set to 1.8 kV, funnel RF level at 45, and heated capillary temperature at 275 °C. The full MS resolution was set to 60,000 at m/z 200 and full MS AGC target was 300% with the injection time set to Auto. Mass range was set to 350–1500. For fragmentation spectra, a resolution of 15,000, Isolation width was set at 1.6 m/z, normalized collision energy was set at 30%. The AGC target value was to Standard, with max injection time of 40msec and we did TopN=30scans.

For Label-Free quantitation search was carried out on the raw files via FragPipe v.19. (Nesvizhskii, A. I. lab, University of Michigan). Data were searched against reviewed Uniprot FASTA for mus musculus and homo sapiens were downloaded from Uniprot and a database of 112 common laboratory contaminants (https://www.thegpm.org/crap/) were used. Precursor mass tolerance was set to -/+ 20ppm and 20ppm for fragments. Trypsin was specified as protease, and a maximum of two missed cleavages was allowed, the required minimum peptide sequence length was 7 amino acids, and the peptide mass was between 500-5000 Da. Carbamidomethylation on cysteine (+57.021 Da) was set as static modification. Dynamic modifications included protein N-terminal acetylation (+42.01Da), oxidation on methionine (+15.995 Da) and deamidation of asparagine and glutamine (0.985 Da). To determine and control the number of false-positive identifications, MsFragger applies a target-decoy search strategy. The false discovery rate (FDR) at less than 1% for peptide spectrum matches and protein group identifications. Label-free protein quantification was performed with the IonQuant algorithm, v.8.10, and *Match Between Runs* feature (MBR) was enabled. We accepted identifications with at least one unique peptide. Table S7 contains extra details relevant data dependent analysis.

#### Data Independent Analysis

Data independent analysis was used for the following samples: immunoprecipitation of nuclear enriched lysate from PND2 *Chd8^+/+^* mouse forebrain, immunoprecipitation of nuclear enriched lysate from PND2 *Chd8^5bpdel/+^* mouse forebrain, and TurboID pulldown of whole cell lysate from HEK293T cells. Peptides were directly eluted onto an Orbitrap Exploris 480 instrument (Thermo Fisher Scientific, Bremen, Germany). Here, spectra were obtained using data-independent analysis (DIA). Spray voltage were set to 1.8 kV, funnel RF level at 45, and heated capillary temperature at 275 °C. For full MS, resolution was set to 120,000 at m/z 200 and full MS AGC target was 300% with an IT of 45msec. Mass range was set to 350–1400. AGC target value for fragment spectra was set at 1000. For the fragmentation spectra 34 windows of 21Da were used with an overlap of 1 Da. Resolution was set to 15,000 and IT=23msec. Normalized collision energy was set to 30%. All data were acquired in profile mode using positive polarity and peptide match was set to off, and isotope exclusion was on.

LCMS files were processed with Spectronaut v.18 (Biognosys, Zurich, Switzerland) using DirectDIA analysis mode. Mass tolerance/accuracy for precursor and fragment identification was set to default settings. The reviewed FASTA for mus musculus and homo sapiens were downloaded from Uniprot and a database of 112 common laboratory contaminants (https://www.thegpm.org/crap/) were used. A maximum of two missing cleavages were allowed, the required minimum peptide sequence length was 7 amino acids, and the peptide mass was limited to a maximum of 4600 Da. Carbamidomethylation of cysteine residues was set as a fixed modification, and methionine oxidation and acetylation of protein N termini as variable modifications. A decoy false discovery rate (FDR) at less than 1% for peptide spectrum matches and protein group identifications was used for spectra filtering (Spectronaut default). Decoy database hits, proteins identified as potential contaminants, and proteins identified exclusively by one site modification were excluded from further analysis. Table S7 contains extra details relevant data independent analysis.

#### PPI scoring and network visualization

Protein interactions were scored with both Significance Analysis of INTeractome Express (SAINTexpress)^58,59^ and Differential Enrichment analysis of Proteomics data (DEP)^60,61^ platforms. For SAINTexpress scoring, the Galaxy APOSTL server under default parameters. IgG bait served as the source of control counts for all IP-MS experiments and CHD8-Turbo -Biotin as the source of control counts for the TurboID experiment. Concurrently, statistical analysis of protein interactions was performed using the R package DEP. Briefly, preprocessed proteomics data was log2-transformed and filtered such that proteins missing from no more than one replicate in at least one group were kept. The data was normalized using variance stabilized transformation. Remaining missing values were imputed using the “MinProb” function within DEP. All PPI networks were built using STRING in Cytoscape (v3.10.3). GO term enrichment was calculated via STRING Enrichment, with annotations for enriched GO terms manually curated.

### Western blot analysis

Samples were diluted in 30μl 6X Laemmli SDS buffer (375mM Tris-HCl, 9% SDS, 50% glycerol, 0.03% Bromophenol blue) and 5% β-mercaptoethanol, boiled at 70℃ for 10min, and separated on 4-20% polyacrylamide tris-glycine protein gel (BioRad). Instead of 70℃ for 10min, samples eluted from beads were boiled at 95℃ for 5min before being separated on the polyacrylamide gel. The separated proteins were transferred onto a PVDF membrane (Millipore Sigma) by wet transfer (overnight, 13mA, 4℃).

Membranes were blocked with Intercept PBS blocking buffer (Li-Cor) at room temperature for one hour.

Primary antibodies were diluted in 7.5ml Intercept PBS blocking buffer with 0.1% Tween. Membranes were incubated with the primary antibody solution overnight at 4℃, then washed four times for 10min with PBS with 0.1% Tween (PBST). Fluorescently tagged secondary antibodies (Li-Cor) were diluted in 10ml Intercept PBS blocking buffer with 0.1% Tween. After the initial washes, blots were incubated with the secondary antibody solution for one hour at room temperature. Blots were washed an additional four times for 10min with PBST and two times with PBS. Bands were visualized using the Odyssey DLx imaging system (Li-Cor).

### Immunohistochemistry

IHC was performed on free-floating sections from PND2 mouse brains according to established protocols in the lab. In short, brains were fixed in 4% paraformaldehyde, cryoprotected in sucrose, and sectioned at 30-40 µm using a cryostat. Sections were stored in PBS with sodium azide until use.

Antigen retrieval was carried out by incubating sections in 1× citrate buffer at 60 °C for 1 hour (Sigma- Aldrich, Cat# C9999). Sections were then washed 3 times in PBS containing 0.1% Triton X-100 (PBST) for 10 minutes each, followed by permeabilization with PBS containing 0.5% Triton X-100 for 20 minutes at room temperature. Blocking was performed in 5% skimmed milk dissolved in 0.1% PBST for 1 hour at room temperature or overnight at 4 °C on a shaker. Primary antibodies were diluted in centrifuged blocking buffer and incubated overnight at 4 °C on a shaker. The next day, sections were washed 3 times in PBST (20 minutes each), then incubated with AlexaFluor-conjugated secondary antibodies (1:1000 in blocking buffer) for 1 hour at room temperature, protected from light. Sections were washed again (3 × 20 minutes in PBST), stained with Hoechst (1:10000 in PBS) for 30 minutes, followed by a final PBS wash. Sections were transferred to glass slides using a paintbrush, allowed to air dry briefly, and mounted using ProLong Gold Antifade Mountant (Invitrogen). For ICC, HEK293T cells or primary neurons were plated on poly-L-lysine-coated coverslips, fixed with 4% paraformaldehyde, permeabilized in 0.5% Triton X-100, and blocked in 5% BSA with 5% goat serum.

Cells were incubated with primary antibodies overnight at 4°C, washed, and stained with fluorescently conjugated secondary antibodies. Nuclei were counterstained with Hoechst and coverslips were mounted using ProLong Gold for imaging.

### Proximity Ligation Assay

Proximity ligation assays were performed using the Duolink® In Situ Red Starter Kit Mouse/Rabbit (Sigma-Aldrich, Cat# DUO92101), following the manufacturer’s protocol with minor adjustments. Cells were seeded on poly-L-lysine-coated coverslips, fixed with 4% paraformaldehyde, and permeabilized using 0.5% Triton X-100 in PBS. Samples were then blocked with Duolink® Blocking Solution for 1 hour at 37 °C in a humidity chamber. After blocking, samples were incubated with primary antibodies raised in rabbit and mouse, diluted in Duolink® Antibody Diluent, and incubated overnight at 4 °C in a humidified chamber. Following two washes in Buffer A (provided with the kit), PLA PLUS and MINUS probes were diluted 1:5 in Antibody Diluent and applied to the samples for 1 hour at 37 °C. After two additional Buffer A washes, ligation was performed using the Duolink® Ligation Solution freshly prepared with Ligase (1:40 dilution), and incubated for 30 minutes at 37 °C. Slides were then washed and incubated in freshly prepared Amplification Solution containing Polymerase (1:80 dilution) for 100 minutes at 37 °C in the dark. Final washes were performed with Buffer B (2 × 10 minutes) followed by a 1-minute wash in 0.01× Buffer B. Slides were mounted using Duolink® In Situ Mounting Medium with DAPI and imaged by fluorescence microscopy using a 20x or 40x objective. PLA signals were visualized as discrete fluorescent puncta indicating close proximity (<40 nm) of target proteins.

### RNA Immunoprecipitation

Immunoprecipitation of nuclear lysate was performed as described above up to the elution. Washed beads were resuspended in 100μl of RIP buffer (50mM HEPES ph7.5 (Gibco 15630106), 100mM NaCl, 5mM EDTA, 10mM DTT, 0.5% TritonX-100, 10% Glycerol, 1% SDS). To determine if RNA is present after immunoprecipitation, RNA was eluted via treatment with proteinase k (Thermo Scientific 25530049). Concentration of RNA was quantified using the Qubit RNA High Sensitivity Assay kit (Invitrogen Q32852) with the Qubit 4 Fluorometer (Invitrogen Q33238) according to the manufacturer’s directions. For nuclease treated samples, RNA was treated with 3μg of RNaseA (Thermo Scientific EN0531) or with 2ul of DNase I (Thermo Scientific EN0521) for 30 minutes at 37°C before quantification.

For RNA immunoprecipitation sequencing (RIP-seq), after resuspension in RIP buffer 500μl of TRIzol (Invitrogen 15596026) was applied to beads still coupled to protein complexes. Beads were incubated for 5 minutes to ensure dissociation of the nucleoprotein complex. Beads were pelleted using a magnetic rack and the supernatant was collected. RNA was purified using a Direct-zol RNA Microprep (Zymo R2060) according to the manufacturer’s directions. Concentration of RNA was quantified using the Qubit RNA High Sensitivity Assay kit (Invitrogen Q32852) with the Qubit 4 Fluorometer (Invitrogen Q33238) according to the manufacturer’s directions. Purified RNA was sent to Novogene for sequencing.

### RIP-Sequencing and bioinformatics analysis

RNA libraries were prepared at Novogene using Illumina reagents. Libraries were sequenced using Illumina NovaSeq 6000 S4 system, paired-end 150 (PE150) method. Reads were aligned to mouse genome (GRCm38/mm10) using STAR (version 2.5.4b)^104^, and gene counts were produced using featureCounts^105^. Data quality was assessed using FastQC^106^, and principal component analysis (PCA) was used to determine presence of sample outliers, all 12 samples passed QC. Bioinformatic analysis was performed using R programming language version 4.4.2 (R Development Core Team, 2015) and RStudio integrated development environment version 2024.12.1 (Team R, 2018). Plots were generated using ggplot2 R package version 3.5.2. Information on sample usage and replicate relevant to RIP- Sequencing can be found in Table S7.

### RIP-seq differential expression (DE) analysis

For DE analysis we used DESeq2 R package^107^. Prior to DE analysis aligned count matrices were filtered to contain at least 20 gene-body counts in a minimum of 1 IP sample to remove ambient RNA and ensure out expression data was the result of IP pull down. This was done individually for each antibody comparison to IgG. DE analysis was performed comparing the antibody of interest to the IgG control group. DESeq2 size-factor normalization, accounting for sequencing depth and RNA composition, was used to generate normalized count matrices that were used for plotting summary heatmaps and expression data of individual genes. PCA was performed using the top 500 variable genes from variance stabilized counts output in DESeq2.

### Gene ontology enrichment analysis

To test for enrichment of GO terms we used the clusterProfiler R package version 4.14.6^108^. Mouse Gene Ontology (GO) data was downloaded from Bioconductor (org.Mm.eg.db). For the analysis presented here, we tested for GO Biological Process, Cellular Component, and Molecular Function annotations. We used a minimal node size of 3 and max of 2000 in order to test against a comprehensive set of annotations, but only considered terms with at least 5 significantly DE genes in a GO term. clusterProfiler uses a Benjamini-Hochberg false discovery rate correction of p-values, which we used to rank enriched gene sets used for plotting. For RNA-seq analysis, we reported terms with FDR < 0.1. The test set of DE genes was compared against the background set of genes expressed in our study based on criteria used for DEG detection, protein coding and an average of at least 100 normalized counts across samples. For RIP-seq analysis, we reported terms with FDR < 0.2. Because we are testing for IP enrichment, the test set of DE genes was compared against the background set of genes that was used for initial alignment.

### RIP-seq permutation testing

To determine significance of the overlap between genes identified in CHD8 CHIP-seq and TaDa-seq and those enriched in Chd8C RIP-seq, we used a permutation test. We used the top 1000 genes enriched in the Chd8C vs IgG DE set by pvalue as our RIP-seq set. Chd8 CHIP-seq data was obtained from the metanalysis performed in (Wade 2019), significant hits in each sample had a “score” greater than 0.1. Due to differences in sensitivity across CHIP-seq experiments the criteria for inclusion differed in the Gompers and Platt sets. For the Gompers set we included any gene that had a “score” greater than 0.1 in one sample (n= 2) which resulted in 856 genes and the Platt we only considered genes with “score” greater than 0.1 in every sample (n= 4) which resulted in 4,582 genes. For the TaDa-seq data we restricted to promotor enrichment resulting in 3,796 genes. Due to the size of the TaDa-seq and Platt CHIP-seq sets we down sampled to the top 1000 genes by enrichment, full sets are shown in the supplementary. To generate a null distribution, we performed 10,000 permutations. In each iteration, a random set of genes equal in size to the RIP list was sampled without replacement from the background gene set, defined as all genes with at least 20 genebody counts in either the IgG or Chd8C samples. The overlap between the CHIP/TaDa set and each permuted gene set was recorded. And a p- value was calculated as the proportion of permutations in which the overlap was greater than or equal to the observed overlap.

## Resource availability

### Lead contact

Requests for further information and resources should be directed to and will be fulfilled by the lead contact, Alex Nord.

### Materials availability

All plasmids generated in this study in this study are available from the lead contact without restriction.

### Data and code availability

The mass spectrometry proteomics data have been deposited to the ProteomeXchange Consortium via the PRIDE^109^ partner repository with identifier PXD070948. RIP-seq datasets are available via the GEO repository (GSE311408). Any additional information required to reanalyze the data reported in this paper is available from the lead contact upon request.

