## Supplementary_Figures for "Molecular interactions of Chd8 in mouse brain highlights a role in chromatin-associated RNA processing"

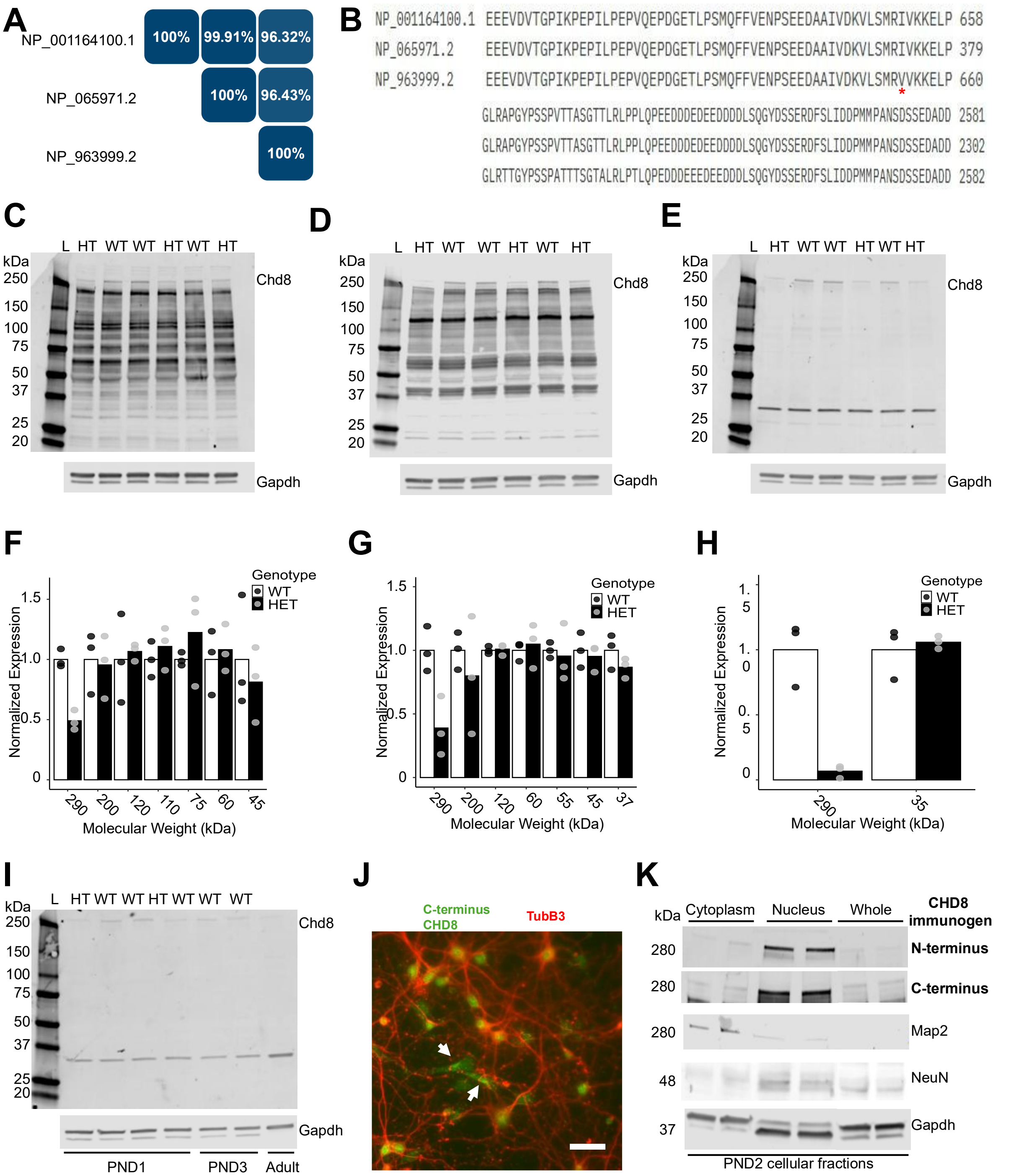


**Figure S1. Validation of Chd8 antibodies and protein localization A-B)** Human and mouse CHD8 protein sequences were downloaded from the NCBI database and aligned using the UniProt Align Tool. NP_001164100.1 and NP_065971.2 are human isoforms and NP_963999.2 is the mouse isoform. **A)** Percent identity matrix of the full CHD8 protein sequences across species. **B)** Alignment of the antibody epitopes across species. **C-E)** Western blotting of PND2 *Chd8*$^{+/+}$ and *Chd8*$^{5bpdel/+}$ mice using the N-terminus of C-terminus Chd8 antibody. Gapdh included as a loading control (Sigma-Aldrich, G8795). **C)** Bethyl N-terminus (A301-224A). **D)**

Abcam N-terminus (ab114126). **E)** Bethyl C-terminus (A301-225A). **F-H)** Quantification of major bands observed in the Chd8 western blots above. Quantification was performed via ImageJ and Chd8 expression was normalized to expression of Gapdh. **F)** Bethyl N-terminus (A301-224A). **G)** Abcam N-terminus (ab114126). **H)** Bethyl C-terminus (A301-225A). **I)** Western blotting of PND1, PND2, and adult *Chd8*$^{+/+}$ and *Chd8*$^{5bpdel/+}$ mice using the C-terminus Chd8 antibody from Abcam (ab224830). **J)** Immunocytochemistry of CHD8 localization in DIV14 cultures co-labeled with the neuronal marker Tubb3. White arrows indicate extranuclear Chd8 signal observed in some Tubb3-negative cells (Scale bar = 50μm). **K)** Western blotting of cytoplasmic, nuclear, and whole cell protein lysates from PND2 mouse forebrain. (N-terminus: Bethyl, A301-224A, C-terminus: Bethyl, A301-225A, Map2: Invitrogen, 13-1500, NeuN: Millipore, MAB377, Gapdh: Sigma-Aldrich, G8795)

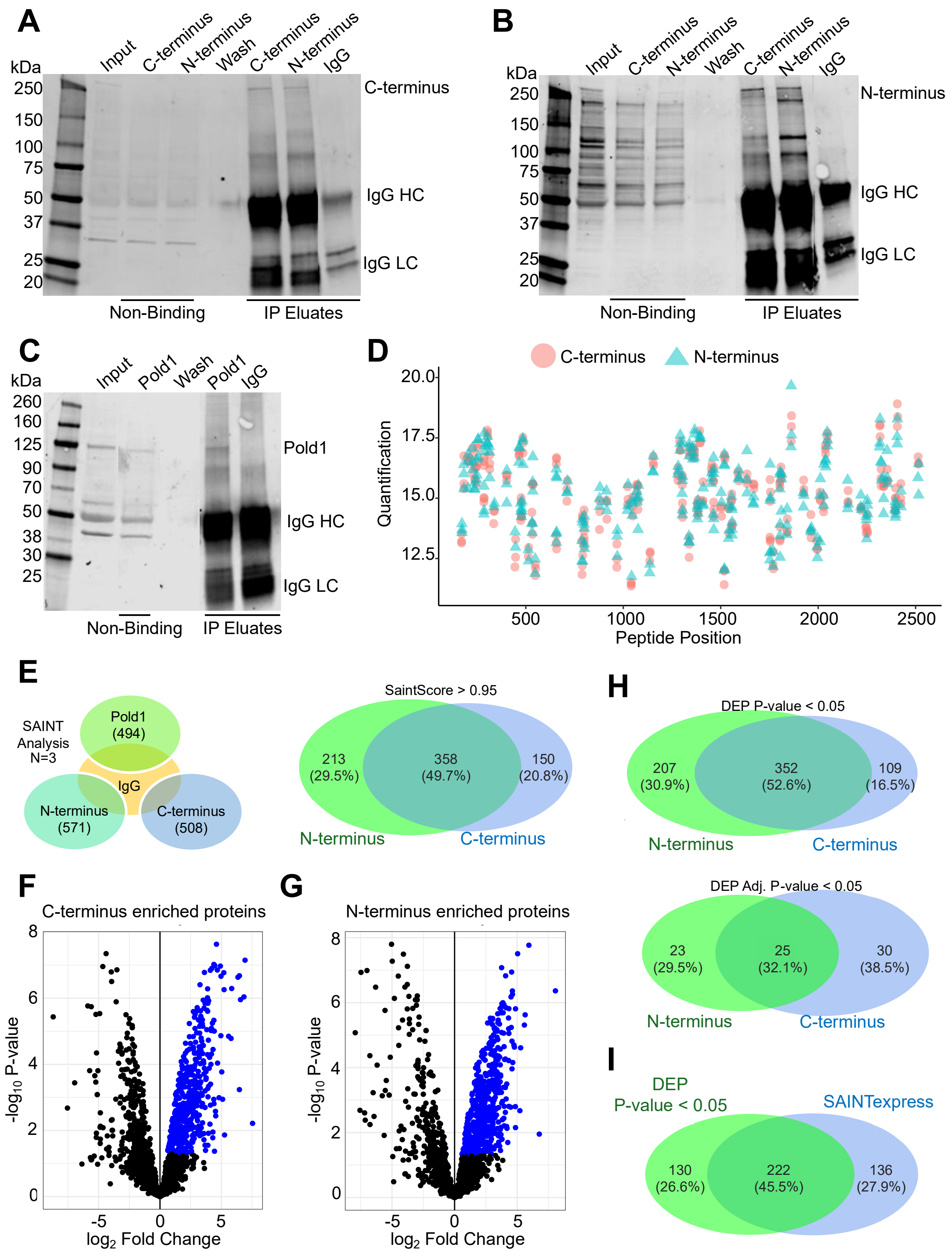

A
kDa
250
150
100
75
50
37
25
20
Input
C-terminus
N-terminus
Wash
C-terminus
N-terminus
IgG
C-terminus
IgG HC
IgG LC
Non-Binding
IP Eluates
B
N-terminus
C
kDa
260
160
125
90
70
50
38
30
25
Input
Pold1
Wash
Pold1
IgG
Pold1
D
C-terminus
N-terminus
Quantification
Peptide Position
E
SAINT Analysis N=3
Pold1 (494)
IgG
N-terminus (571)
C-terminus (508)
SaintScore > 0.95
213 (29.5%)
358 (49.7%)
150 (20.8%)
H
DEP P-value < 0.05
207 (30.9%)
352 (52.6%)
109 (16.5%)
DEP Adj. P-value < 0.05
23 (29.5%)
25 (32.1%)
30 (38.5%)
I
DEP P-value < 0.05
SAINTexpress
130 (26.6%)
222 (45.5%)
136 (27.9%)
F
C-terminus enriched proteins
G
N-terminus enriched proteins
-log10 P-value
log2 Fold Change

**Figure S2. Endogenous IP from mouse brain and Chd8 PPI scoring. A-C)** Western blot validation of successful IP using the Chd8 or Pold1 antibodies. Labels on the top indicate which antibody was used for IP. Labels on the right indicate the antibody used for blotting and the protein band of interest. IgG heavy chain and light chain contamination also indicated on the right. **A)** Chd8 IP with C-terminus blotting. **B)** Chd8 IP with N-terminus blotting. **C)** Pold1 IP with Pold1 blotting. **D)** Peptide coverage map of Chd8 after IP using either the C-terminus or N-terminus Chd8 antibody. **E)** SAINTexpress scoring of PPIs against IgG. Proteins with a SaintScore > 0.95 for either the C-terminus or N-terminus antibody were identified and then compared. **F-G)** Volcano plots of DEP scoring of PPIs against IGG. Blue dots indicate proteins that are enriched in C-terminus or N-terminus IP compared to IgG controls that pass a significance cutoff of P-value < 0.05. **F)** C-terminus. **G)** N-terminus. **H)** Comparison of proteins that were identified by DEP in C-terminus or N-terminus IP, using significance cutoffs of P-value < 0.05 or Adjusted P-value < 0.05. **I)** Comparison of the two PPI scoring methods as the final filtering step to determine Chd8 protein interactors.

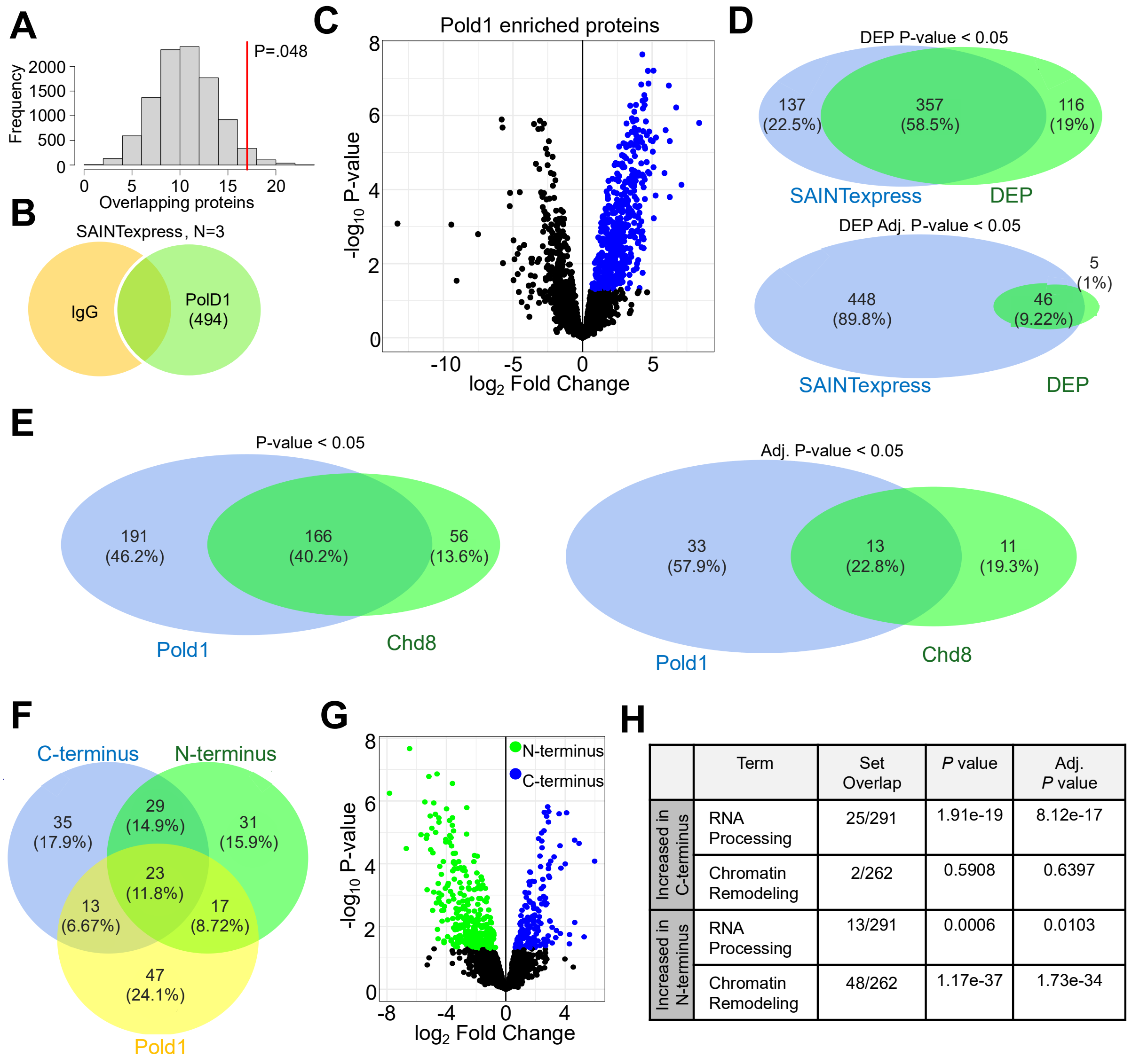


| | Term | Set Overlap | *P* value | Adj. *P* value |
|---|---|---|---|---|
| Increased in C-terminus | RNA Processing | 25/291 | 1.91e-19 | 8.12e-17 |
| | Chromatin Remodeling | 2/262 | 0.5908 | 0.6397 |
| Increased in N-terminus | RNA Processing | 13/291 | 0.0006 | 0.0103 |
| | Chromatin Remodeling | 48/262 | 1.17e-37 | 1.73e-34 |

**Figure S3. Pold1 PPI scoring and hd8 antibody comparison A)** Permutation test of enrichment in high-confidence SFARI genes (Category 1). All proteins detected by mass spectrometry after Chd8 IP and all listed SFARI genes were used as background. P=0.48 **B)** SAINTexpress scoring of PPIs against IgG. Pold1 enriched proteins were identified by having a SaintScore > 0.95. **C)** Volcano plot of DEP scoring of PPIs against IgG. Blue dots indicate proteins that are enriched in Pold1 IP compared to IgG controls that pass a significance cutoff of P-value < 0.05. **D)** Comparison of the two PPI scoring methods using either P-value < 0.05 or Adjusted P-value < 0.05. **E)** Comparison Chd8 and Pold1 enriched PPIs. Proteins were identified by SAINTexpress and DEP at a significance cutoff of either P value < 0.05 or Adjusted P value < 0.05. Chd8 PPIs must have been identified in both C-terminus and N-terminus IP. **F)** Comparison of the top 100 PPIs identified in C-terminus, N-terminus, and Pold1 IPs. PPIs were ordered by significance via P-value. **G)** Volcano plot of a DEP analysis comparing the Chd8 antibodies. Green dots indicate proteins enriched in the N-terminus pulldown and blue dots indicate proteins enriched in the C-terminus pulldown, with a significance cutoff of P < 0.05. **H)**

GO term enrichment of the proteins significantly enriched in either the N-terminus or C-terminus pulldowns. GO term enrichment performed with Enrichr with default settings.

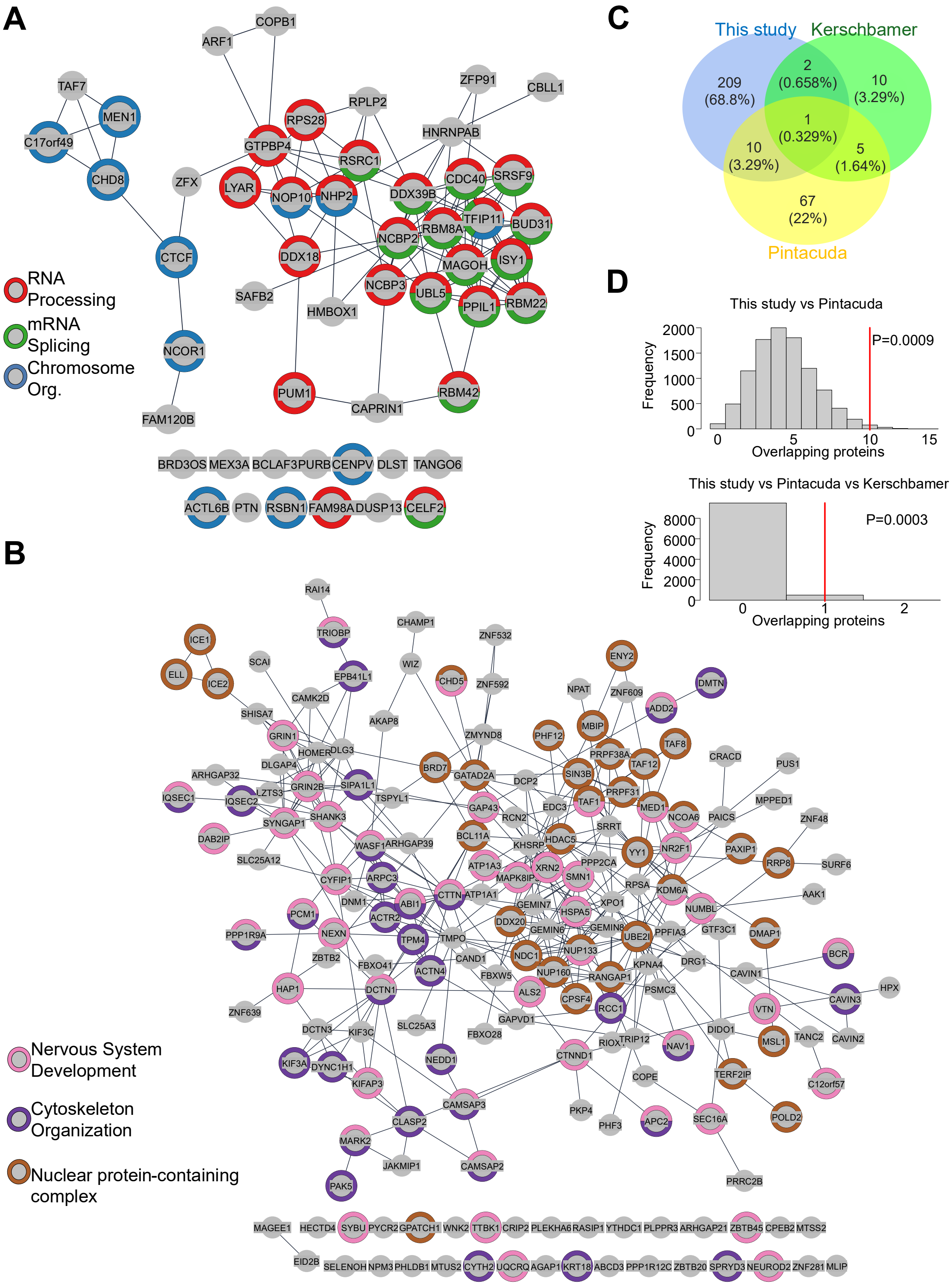
A
COPB1
ARF1
TAF7
MEN1
C17orf49
CHD8
ZFX
CTCF
NCOR1
FAM120B
RPLP2
RPS28
GTPBP4
RSRC1
LYAR
NOP10
NHP2
DDX39B
CDC40
SRSF9
ZFP91
CBLL1
HNRNPAB
TFIP11
BUD31
RBM8A
NCBP2
DDX18
MAGOH
ISY1
NCBP3
UBL5
PPIL1
RBM22
SAFB2
HMBOX1
PUM1
CAPRIN1
RBM42
BRD3OS
MEX3A
BCLAF3
PURB
CENPV
DLST
TANGO6
ACTL6B
PTN
RSBN1
FAM98A
DUSP13
CELF2
RNA Processing
mRNA Splicing
Chromosome Org.
B
Nervous System Development
Cytoskeleton Organization
Nuclear protein-containing complex
RAI14
TRIOBP
CHAMP1
ZNF532
ICE1
ELL
ICE2
SCAI
EPB41L1
WIZ
CHD5
ZNF592
ENY2
NPAT
ZNF609
DMTN
ADD2
CAMK2D
SHISA7
AKAP8
ZMYND8
PHF12
MBIP
TAF8
GRIN1
HOMER
DLG3
DLGAP4
BRD7
GATAD2A
PRPF38A
TAF12
CRACD
PUS1
ARHGAP32
SIPA1L1
GRIN2B
LZTS3
IQSEC1
IQSEC2
TSPYL1
DCP2
SIN3B
PRPF31
MPPED1
GAP43
EDC3
TAF1
MED1
NCOA6
PAICS
ZNF48
SHANK3
SYNGAP1
RCN2
SRRT
DAB2IP
WASF1
ARHGAP39
BCL11A
KHSRP
HDAC5
YY1
NR2F1
PAXIP1
RRP8
SURF6
SLC25A12
ATP1A3
PPP2CA
XRN2
CYFIP1
ARPC3
MAPK8IP
SMN1
RPSA
KDM6A
AAK1
DNM1
ABI1
CTTN
ATP1A1
GEMIN7
XPO1
NUMBL
PCM1
ACTR2
HSPA5
DDX20
GEMIN8
PPP1R9A
NEXN
TPM4
TMPO
GEMIN6
UBE2I
PPFIA3
GTF3C1
DMAP1
ZBTB2
FBXO41
CAND1
NDC1
NUP133
KPNA4
DRG1
BCR
HAP1
DCTN1
ACTN4
FBXW5
NUP160
RANGAP1
CAVIN1
HPX
ALS2
CPSF4
PSMC3
CAVIN3
ZNF639
RCC1
VTN
SLC25A3
GAPVD1
DIDO1
DCTN3
KIF3C
FBXO28
RIOX1
TRIP12
NAV1
MSL1
TANC2
CAVIN2
KIF3A
DYNC1H1
NEDD1
CTNND1
COPE
TERF2IP
C12orf57
KIFAP3
CAMSAP3
PKP4
SEC16A
CLASP2
PHF3
APC2
POLD2
MARK2
JAKMIP1
CAMSAP2
PRRC2B
PAK5
MAGEE1
HECTD4
SYBU
PYCR2
GPATCH1
WNK2
TTBK1
CRIP2
PLEKHA6
RASIP1
YTHDC1
PLPPR3
ARHGAP21
ZBTB45
CPEB2
MTSS2
EID2B
SELENOH
NPM3
PHLDB1
MTUS2
CYTH2
UQCRQ
AGAP1
KRT18
ABCD3
PPP1R12C
ZBTB20
SPRYD3
NEUROD2
ZNF281
MLIP
C
This study
Kerschbamer
Pintacuda
209
(68.8%)
2
(0.658%)
10
(3.29%)
1
(0.329%)
10
(3.29%)
5
(1.64%)
67
(22%)
D
This study vs Pintacuda
Frequency
2000
1500
1000
500
0
0
5
10
15
Overlapping proteins
P=0.0009
This study vs Pintacuda vs Kerschbamer
Frequency
8000
6000
4000
2000
0
0
1
2
Overlapping proteins
P=0.0003

**Figure S4. Chd8 and Pold1 exclusive PPIs and comparison with previously published Chd8 PPI data. A-B)** STRING network of PPIs exclusive to Chd8 or Pold1. Proteins must have been identified by SAINTexpress and DEP at P value < 0.05, with Chd8 proteins identified by both C- and N-terminus antibodies. Color of the outer ring indicates GO term annotation. Networks were made using default settings for full STRING network indicating both functional and physical protein associations. **A)** Chd8 exclusive proteins. **B)** Pold1 exclusive proteins. **C)** Comparison of Chd8 PPIs identified in this study compared to previously published Chd8 proteomic data. Chd8 proteins included from this study must have been identified by both C- and N-terminus antibodies using SAINTexpress and DEP at P value < 0.05. **D)** Permutation test of the overlap in PPIs identified across published Chd8 proteomics data. Overlaps between this study and the Pintacuda publication and overlaps between all three publications were significant (P=0.0009 and P=0.0003, respectively). The overlap between this study and the Kerschbamer publication was not significant. All proteins detected in mass spectrometry after any IP in this or other studies were used as background.

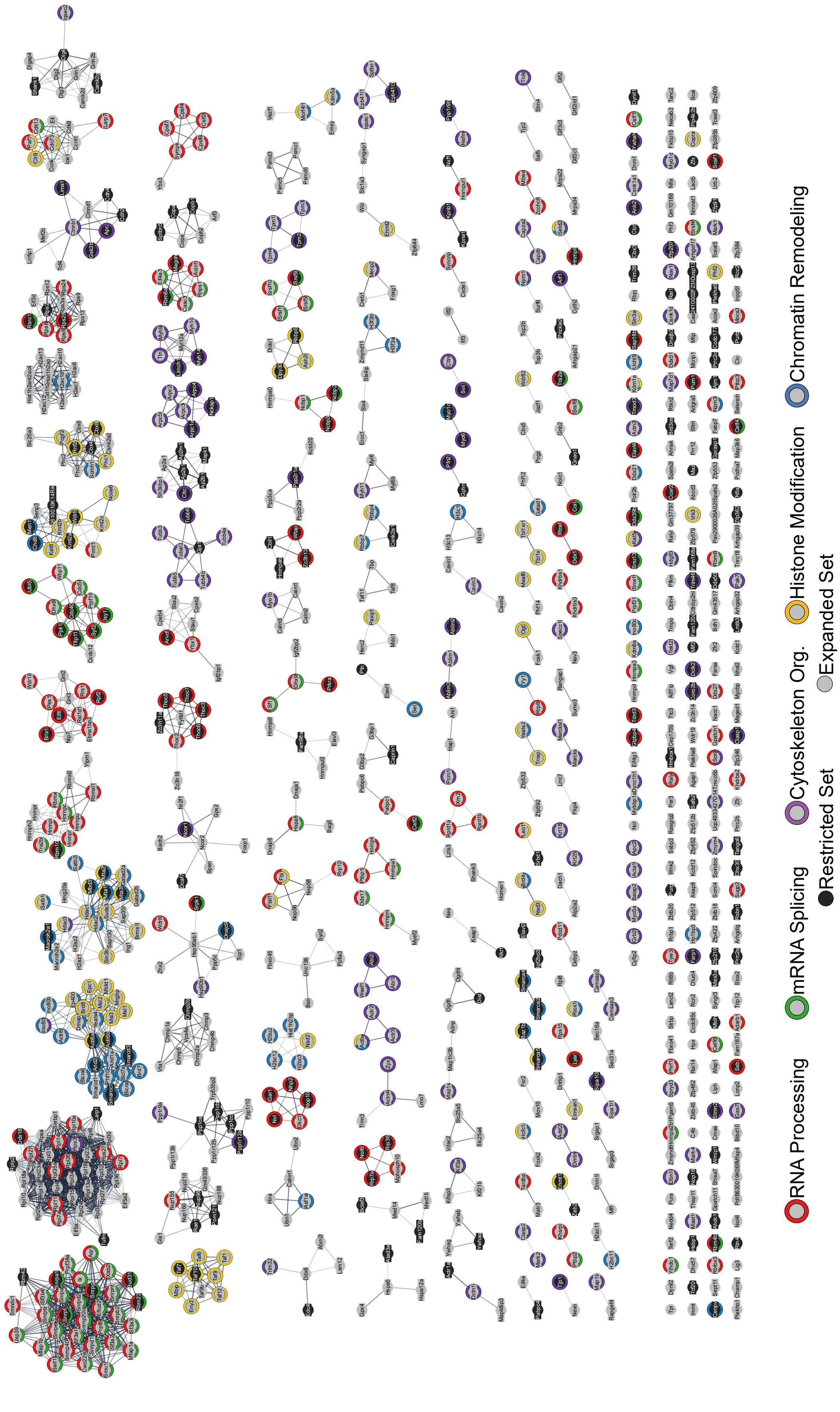
RNA Processing
mRNA Splicing
Cytoskeleton Org.
Histone Modification
Chromatin Remodeling
Restricted Set
Expanded Set

**Figure S5. Full MCL clustering of expanded Chd8 PPI network.** All clusters associated with the MCL analysis that was highlighted in Figure 2. MCL clustering of STRING Chd8 protein-protein network with all proteins identified by either Chd8 antibody via either SAINTexpress or DEP at a P-value < 0.05 significance cutoff. MCL clustering was performed with the granularity parameter set to 6. Color of the outside ring indicates the annotated GO term. Inner circle color indicates if the protein was included in the more stringent 222 Chd8 interacting protein set. Network was made using default settings for physical STRING network indicating the proteins are part of a physical complex.

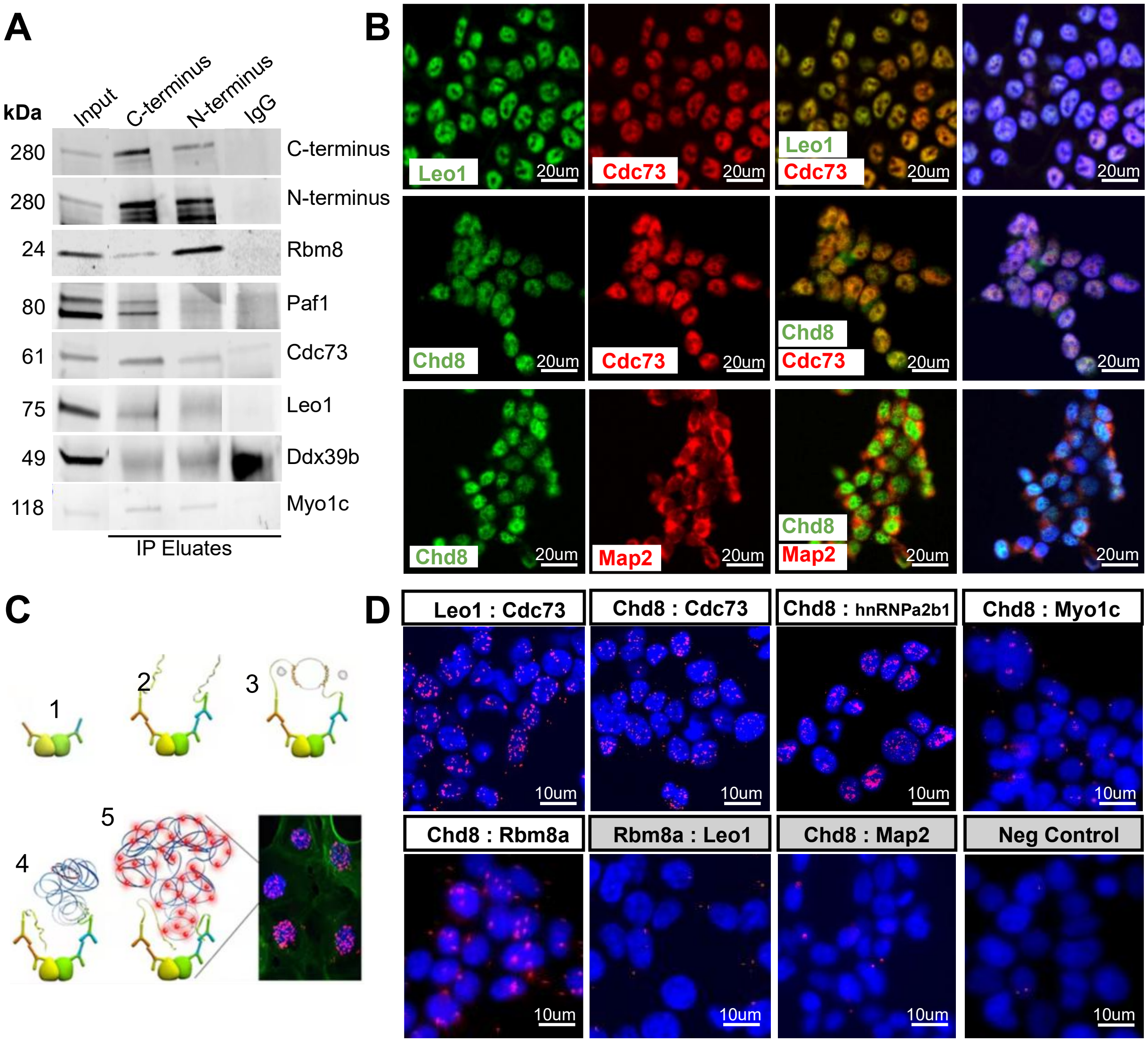


**Figure S6. Validation of Chd8 protein interactions. A)** Western blotting of CoIP validation of Chd8 protein interactions. CoIP was performed using either the C-terminus or N-terminus Chd8 antibody, indicated by top labels. The antibody used for western blotting is indicated to the right and the molecular weight (kDa) of the band to the left. **B)** Colocalization validation of protein interactions. Colocalization of Leo1 and Cdc73 acts as a positive control and colocalization of Chd8 and Map2 acts as a negative control. Far right panel represents a merge with all color channels. Nuclei were stained blue with Hoechst. **C)** Schematic of Proximity Ligation Assay (PLA) approach. **D)** PLA validation of CHD8 protein interactions. Nuclei were stained with Hoechst (blue), and red puncta indicate sites of interaction between the two target proteins. The PLA between LEO1 and CDC73, both members of the PAF complex, serves as a positive control. Gray panels show various negative controls, with the “Neg control” panel representing the no-probe condition.

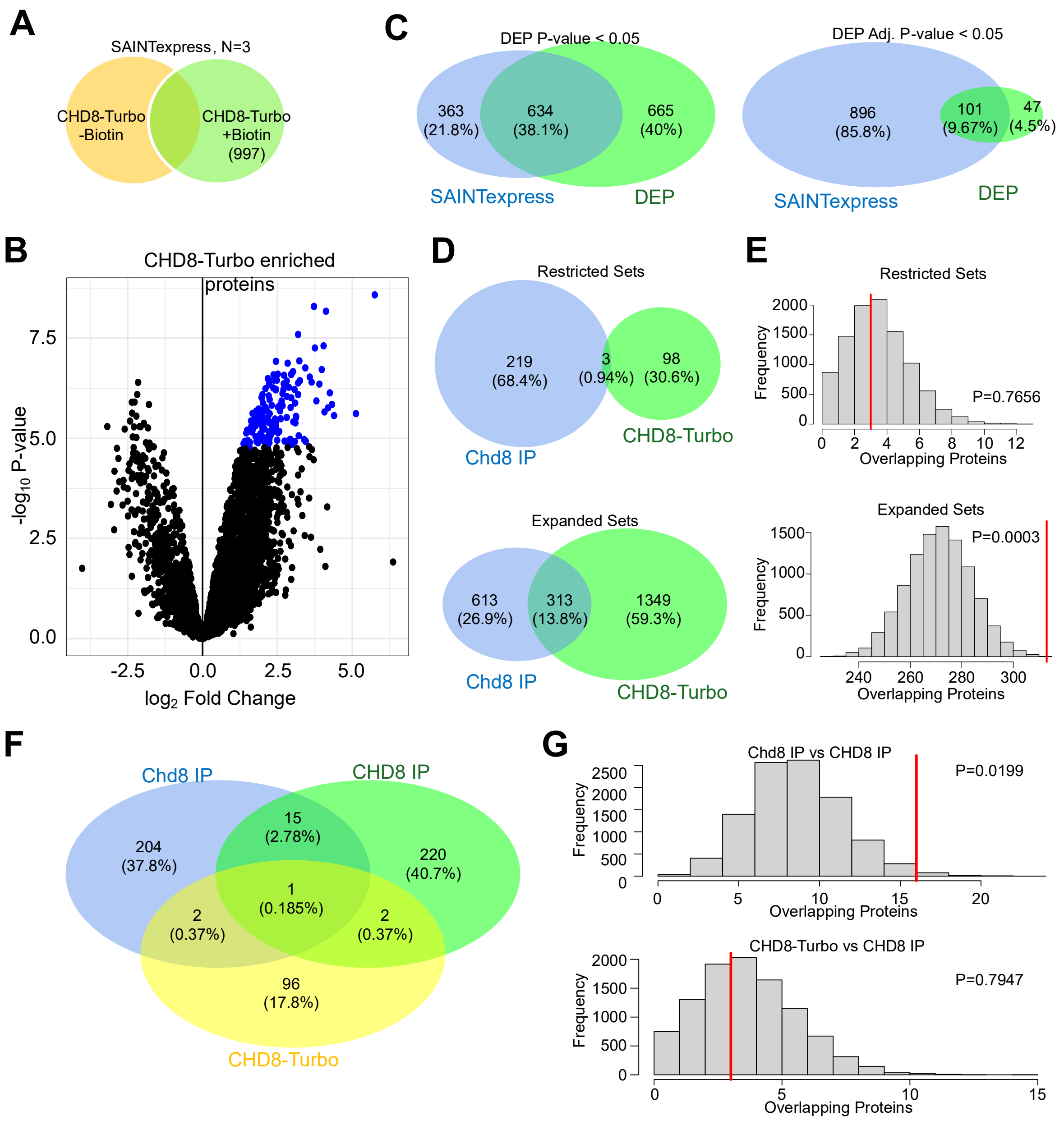


**Figure S7. CHD8-Turbo PPI scoring and comparison to endogenous IP. A)** SAINTexpress PPI scoring of CHD8-Turbo +Biotin compared to CHD8-Turbo -Biotin as a control. Proteins were identified by having a SaintScore > 0.95. **B)** Volcano plot of DEP scoring of PPIs against CHD8-Turbo –Biotin. Blue dots indicate proteins that are enriched in CHD8-Turbo +Biotin compared to CHD8-Turbo -Biotin controls that pass a significance cutoff of Adjusted P value < 0.05. **C)** Comparison of the two PPI scoring methods using DEP at P value < 0.05 or Adjusted P value < 0.05. **D)** Comparison between endogenous Chd8 IP from PND2 mouse forebrain and CHD8-Turbo. Restricted sets include proteins that were significant using both SAINTexpress and DEP, with Chd8 IP at a P value < 0.05 cutoff and TurboID at an adjusted P value < 0.05 cutoff. Chd8 IP proteins must have been significant in both C- and N- terminus proteins. Expanded sets included all proteins that were significant using either SAINTexpress or DEP at a cutoff of P value < 0.05. Chd8 IP proteins could be significant in either C- or n-terminus antibodies. **E)**

Permutation tests of the overlap between endogenous Chd8 IP from PND2 mouse forebrain and CHd8-Turbo with either the restricted or expanded protein sets (P=0.7656 and P=0.0003, respectively). All proteins detected in mass spectrometry after any IP or TurboID condition were used as background. **F)** Comparisons between endogenous Chd8 IP from mouse forebrain, endogenous CHD8 IP from HEK cells, and CHD8-Turbo PPIs. All conditions include proteins that were identified by both SAINTexpress and DEP, with IP conditions including proteins identified in both N- and C-terminus antibodies. **G)** Permutation tests of the overlap between the two IP datasets and between CHD8-Turbo and CHD8 IP in HEK cells. All proteins detected in mass spectrometry after any IP or TurboID condition were used as background.
